# Impacts of increasing isolation and environmental variation on Florida Scrub-Jay demography

**DOI:** 10.1101/2024.01.10.575127

**Authors:** Jeremy Summers, Elissa J. Cosgrove, Reed Bowman, John W. Fitzpatrick, Nancy Chen

## Abstract

Isolation caused by anthropogenic habitat fragmentation can destabilize populations. Populations relying on the inflow of immigrants can face reduced fitness due to inbreeding depression as fewer new individuals arrive. Empirical studies of the demographic consequences of isolation are critical to understand how populations persist through changing conditions. We used a 34-year demographic and environmental dataset from a population of cooperatively-breeding Florida Scrub-Jays (*Aphelocoma coerulescens*) to create mechanistic models linking environmental and demographic factors to population growth rates. We found that the population has not declined despite both declining immigration and increasing inbreeding, owing to a coinciding response in breeder survival. We find evidence of density-dependent immigration, breeder survival, and fecundity, indicating that interactions between vital rates and local density play a role in buffering the population against change. Our study elucidates the impacts of isolation on demography and how long-term stability is maintained via demographic responses.

## Introduction

Identifying the potential demographic and environmental drivers of population dynamics is a fundamental goal of ecology (Cappuccino & Price 1995; Iles *et al*. 2019). Population growth rates are influenced by density-dependent (*e*.*g*., resource competition) and independent (*e*.*g*., climate) factors that interact in ways that remain poorly understood (Goswami *et al*. 2011; Oro 2013). Furthermore, the dynamics of small populations can be affected by intrinsic genetic factors such as inbreeding depression, or decreased fitness of inbred individuals (Fagan & Holmes 2006; Kardos *et al*. 2023). As more populations worldwide are threatened by habitat fragmentation and climate change (Marzluff *et al*. 2001; Simberloff 1999; Young *et al*. 2016), elucidating the effects of temporal changes in immigration, inbreeding, and environmental factors on the demography of wild populations is increasingly urgent (Kardos *et al*. 2023; Schaub & Ullrich 2021).

As populations become more isolated, theory predicts that immigration will decrease and levels of inbreeding will increase, but downstream effects on population dynamics are more complicated to predict. Empirical studies have found that immigration may play an important role in governing population dynamics (Lieury *et al*. 2015; Santoro *et al*. 2016; Tenan *et al*. 2023), but the demographic impact of immigration varies depending on recent changes in metapopulation structure (Reichert *et al*. 2021), resource availability (Horswill *et al*. 2022), temporal variation in the environment (Millon *et al*. 2019), and population density (Harman *et al*. 2020). Decreasing immigration may not always lead to population declines, as immigration may not be a large driver of variability in population growth rates or may be buffered by other demographic and environmental factors. For instance, in classic source-sink systems, source populations that are net producers of immigrants should be insensitive to changes in incoming immigration (Pulliam 1988). Immigration may be negatively density-dependent, which inherently stabilizes populations against fluctuations (Sæther *et al*. 1999; Wilson *et al*. 2017). Finally, temporal variation in immigration could be compensated by changes in other vital rates through density-dependent processes or differential responses to environmental drivers, leading to buffering via demographic compensation (Villellas *et al*. 2015). High local population regulation relative to immigration will make a population more asynchronous from its metapopulation (Lande *et al*. 1999), which will result in greater local stability when the surrounding metapopulation is in decline (Fox *et al*. 2017; Rowland *et al*. 2022). Characterizing the relative contributions of immigration and local survival or fecundity has important implications on the spatial scale required for conservation (Paquet *et al*. 2021), yet few studies have quantified temporal changes in immigration and subsequent effects on population dynamics (Horswill *et al*. 2022).

Gene flow into populations via immigration may counter the negative demographic effects of inbreeding depression (Hedrick & Garcia-Dorado 2016; Niskanen *et al*. 2020; Whitlock *et al*. 2000). The extent of population-level effects of inbreeding depression remains controversial (Bell *et al*. 2019; Keller *et al*. 2007; Kyriazis *et al*. 2020; Mathur & DeWoody 2021; Szulkin *et al*. 2013), and relatively few studies have documented negative impacts of inbreeding on population growth rates (Bozzuto *et al*. 2019; Johnson *et al*. 2011; Kardos *et al*. 2023).Theory predicts that the demographic consequences of inbreeding depression depend on the ecology and life history of a species. Effects of inbreeding on vital rates that do not contribute much to population growth will not result in large changes in population size (Johnson *et al*. 2011; Mills & Smouse 1994). Populations that are primarily regulated by density- or frequency-dependent (soft) selection may not experience reduced population growth in response to inbreeding depression (Saccheri & Hanski 2006). Finally, as the strength of inbreeding depression can vary with environmental conditions, the population-level consequences of inbreeding may also depend on other ecological conditions. More empirical studies that integrate the environmental and demographic contexts of both immigration and inbreeding are needed to clarify the role of increasing isolation on population dynamics.

Quantifying the demographic effects of immigration and inbreeding requires large amounts of data that are rarely available in natural populations. Direct measures of immigration require tracking all new individuals in a population to identify immigrants and residents (Greenwood & Harvey 1982; Hixon *et al*. 2002; Millon *et al*. 2019). While recent advances in population modeling have allowed for the inference of immigration rates in situations where data are limited (Abadi *et al*. 2010), these inferences may overestimate the impact of immigration by failing to account for temporal variation in immigration rates (Nater 2024; Paquet *et al*. 2021). Similarly, investigating the consequences of inbreeding requires measures of relatedness between individuals and individual fitness measures (Johnson *et al*. 2011; Sin *et al*. 2021). Long-term, detailed, population and ecological data obviate the need to infer immigration rates and provide relatedness and fitness data for multiple individuals, and, while rare, provide the best opportunity to empirically test expectations of the consequences of changing immigration rates and inbreeding.

The large amounts of data available through long-term ecological studies can be analyzed with recently developed transient life table response experiments (tLTRE) (Koons *et al*. 2016, 2017). tLTREs retrospectively quantify the contribution of different vital rates to observed variation in population growth rate while accounting for temporal variation in vital rates and demographic structure (Koons *et al*. 2016, 2017). As opposed to prospective sensitivity analyses (Caswell 2000), tLTREs consider both the *potential* impact of vital rates on population growth and the *realized* variation in these rates, making them ideal for attributing observed change in growth rates to underlying change in vital rates (Coulson *et al*. 2005), including immigration (Christian *et al*. 2024). These methods can be built upon to link environmental factors to vital rates through a path analysis (Van Tienderen 2000), ultimately linking variation in the environment to variation in population growth rate (Coulson & Tuljapurkar 2008; Dahlgren & Ehrlén 2009; Knape *et al*. 2023; Nater *et al*. 2021).

Here we empirically test how a natural population of Federally Threatened Florida Scrub-Jays (*Aphelocoma coerulescens*) has remained robust despite increasing isolation. Florida Scrub-Jays are facultative cooperative breeders: most individuals delay dispersal and assist with rearing young on their natal territories, though some disperse and help on other territories before establishing as a breeder (Woolfenden & Fitzpatrick 1984). Females are more likely to disperse earlier and farther than males, but few other vital rates differ between sexes (Aguillon *et al*. 2017; Suh *et al*. 2020; Woolfenden & Fitzpatrick 1984). Florida Scrub-Jays occupy territories year-round and are restricted to fire-mediated xeric oak scrub habitat (Fitzpatrick & Bowman 2016). Habitat loss and degradation has led to declines across their range (Boughton & Bowman 2011; Coulon *et al*. 2008). One color-banded population of Florida Scrub-Jays at Archbold Biological Station has been monitored for >50 years, accumulating complete life histories for thousands of individuals, territory and burn area maps, climate and acorn abundance data, and a 14-generation population pedigree (Chen *et al*. 2019; Fitzpatrick & Bowman 2016). This population has experienced a significant decline in immigration over the past 30 years, which has contributed to increased inbreeding and inbreeding depression (Chen *et al*. 2016), yet census population sizes have not declined (Figure 1).

**Figure 1.**
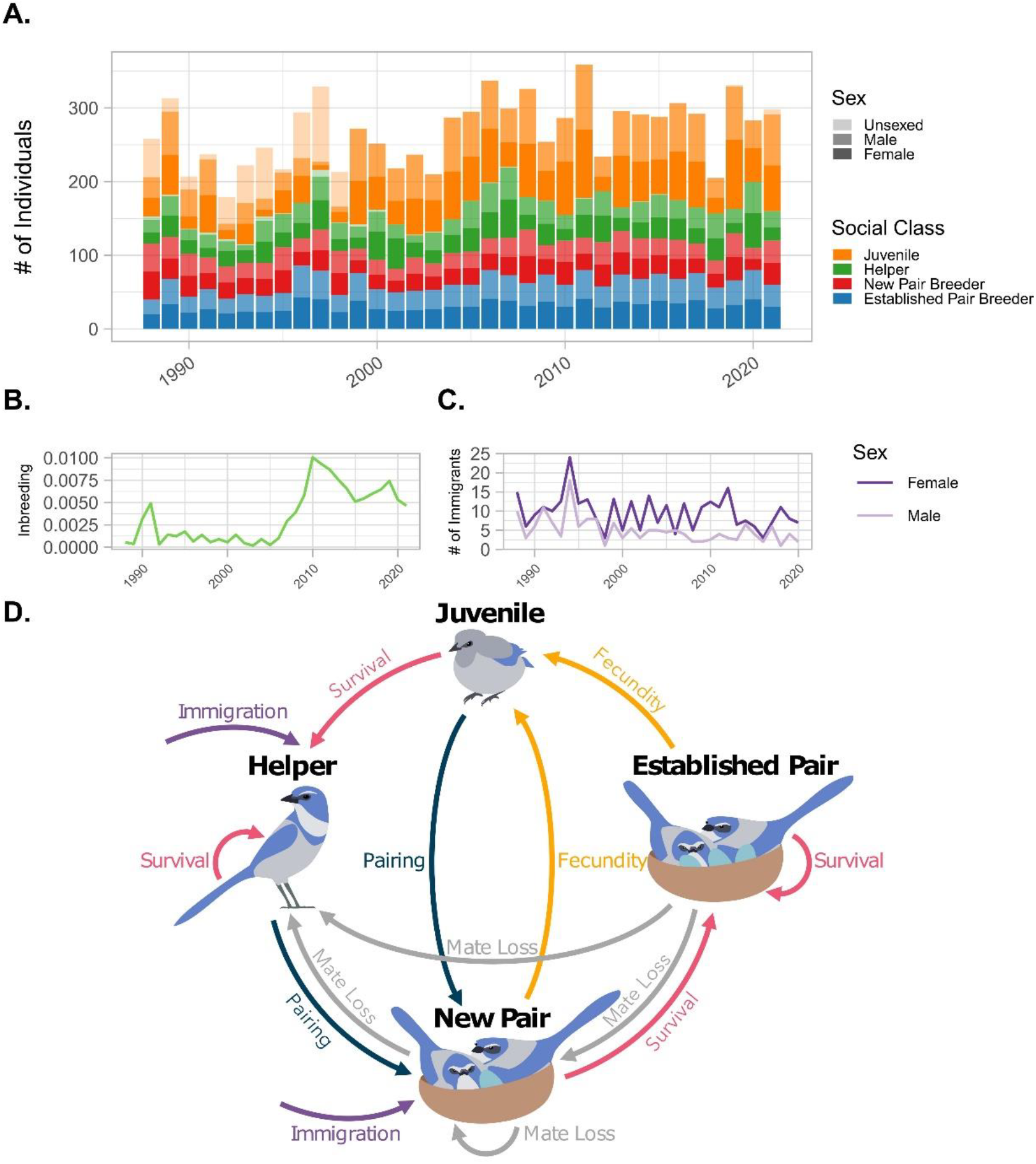
Demography of the Florida Scrub Jay. (A) Total population size over the study period divided by sex and social class. (B) Mean inbreeding coefficient of all individuals over time. (C) Total number of new male and female immigrants arriving to the population over time. (D) Life cycle diagram for the Florida Scrub Jay that describes the movement of individuals across social classes. Individuals who survive to the next breeding season may remain in their existing pair status (pink arrows), form new pairs (dark blue arrows), or lose a mate through divorce or death (gray arrows). New individuals enter the population either through reproduction (orange arrows) or immigration (purple arrows).

We characterized the factors underlying variation in population growth rates over a period of 34 years (1989-2021) to (1) determine the impact of sex-biased immigration on population growth rate, (2) quantify the impact of inbreeding, population density, and other relevant environmental factors on vital rates and population growth rate, and (3) test whether the demographic impacts of decreased immigration and increased inbreeding have been counteracted by other vital rates or environmental factors. We constructed both female-specific and male-specific stage-structured matrix population models and used tLTREs to assess the contribution of different vital rates to population growth. We used generalized linear models (GLMs) to link vital rates to population-wide levels of inbreeding, population density, and environmental factors, and performed environmental tLTREs to determine how these factors have influenced population growth. These analyses allow us to detect density dependence directly and quantify the realized impact of changes in immigration and inbreeding, as well as any other coinciding temporal trend. We predicted that density-dependent responses in the vital rates of non-breeders would compensate for declining immigration, given the abundance of resident non-breeders available to fill breeding vacancies (Woolfenden & Fitzpatrick 1984), and that inbreeding would not strongly influence population growth because of density dependence. Our study is among the few that use direct measures of immigration and inbreeding to determine the impact of isolation on the demography of a natural population, which is crucial for understanding how to preserve threatened populations.

## Methods

### 1. Data Collection

#### 1.1 Study Population

We used data from a long-term study of Florida Scrub-Jays at Archbold Biological Station (Woolfenden & Fitzpatrick 1984). The population is censused monthly, and all nests are tracked during the breeding season (March-June), providing detailed location, survival, and breeding records for all individuals within the study area and allowing identification of immigrants over time. We limited our study area to 14.8 km^2^ of land that has been consistently monitored between 1988-2021. Our dataset includes 4,267 individuals (1,948 female, 1,820 male, and 499 unsexed) from 1,894 territory-years. Please see Supplementary Methods for details on demographic data collection.

#### 1.2 Covariates

We considered several covariates that likely impact Florida Scrub-Jay demography: pedigree-based inbreeding and relatedness coefficients, population density, annual acorn abundance, burned area, and the equatorial southern oscillation index during the dry (breeding) season (EQSOI) (Supplementary Methods). We used mean inbreeding coefficients or pairwise relatedness coefficients estimated from the population pedigree to capture the effect of known inbreeding depression on fecundity and survival in our study population (Chen *et al*. 2016). We calculated population density as the total number of individuals per occupied hectare. As a proxy for habitat quality, we calculated the area burned within the past 2-9 years (the frequency of prescribed fires that generate optimal Florida Scrub-Jay habitat; (Fitzpatrick & Bowman 2016). We used acorn abundance as a metric of resource availability, as acorns are the most important over-winter food source for the Florida Scrub-Jay and vary greatly in abundance over time (DeGange *et al*. 1989; Pesendorfer *et al*. 2021). EQSOI is a commonly used climate index that captures atmospheric conditions that influence El Niño events (Shi & Su 2020), which heavily impact precipitation and temperature in Florida.

### 2. Contribution of Variation in Vital Rates to Population Growth

#### 2.1 Stage-Structured Matrix Population Model

We constructed matrix population models with four stages (juveniles, helpers, new pairs, and established pairs; Figure 1) and a post-breeding census (Supplementary Methods). We defined juveniles as any individuals between 11 days and 1 year of age, which includes all sexually immature individuals. Helpers are individuals of 1 year of age or older who are not established as a socially dominant breeder. Previous work showed that pairs that had bred together previously had higher fecundity than newly-formed pairs (Woolfenden & Fitzpatrick 1984); therefore we divided breeding pairs into new and established pairs based on whether the two individuals had previously paired during a breeding season. Immigrants (all individuals who were not born within the study area) can join the population as either helpers or breeders, forming new pairs. Extensive population monitoring allowed us to fit both female-only and male-only models using direct measures of immigration and sex-specific vital rates for adults. To account for uncertainty caused by unsexed individuals, we randomly assigned these individuals a sex based on a 1:1 sex ratio (Woolfenden & Fitzpatrick 1984) 100 times for each model and calculated annual vital rates for each iteration to parameterize annual projection matrices (Supplementary Methods).

#### 2.2 Transient Life Table Response Experiment

To quantify how vital rates result in observed variation in population growth rates over time, we performed a tLTRE (Koons *et al*. 2016). This method does not assume that the population is near its stable stage distribution and allows variation in stage structure to contribute to variation in population growth rate (Koons *et al*. 2016). We calculated the realized growth rate as the product of the projection matrix (*A(t)*) that is parameterized with the observed vital rates (θ_*i*_, for all vital rates *i*) and the stage distribution vector for each year (*n*). We calculated the sensitivities of the realized growth rate to the vital rates using the *Deriv* package (Clausen *et al*. 2021). We weighted the sensitivity of the population growth rate to each vital rate by the observed variation in vital rates to calculate the variation in growth caused by variation and covariation in vital rates. Covariation can produce a negative contribution when opposite trends in two vital rates result in more similar growth rates over time (*e*.*g*., negative covariation between survival and fecundity means high survival years have low fecundity, reducing fluctuations in population size). To include time-lagged effects, we considered the contribution of variation in vital rates from the same year (*t*) and from the previous year (*t* - 1) to variation in the growth rate. Please see Supplementary Methods for details.

### 3. Impact of Inbreeding, Density, and Environmental Factors on Vital Rates

To determine how environmental variation influences population growth rate, we modeled the underlying vital rates in terms of the covariates described above. We used GLMs with a binomial distribution to model survival for each social class (juveniles, helpers, and breeders) and transition rates from non-breeding to breeding status. We ran separate models for male and female helpers and breeders but not for juveniles due to the large number of unsexed juveniles (480). Note that because Florida Scrub-Jays typically disperse as helpers (Suh *et al*. 2020; Woolfenden & Fitzpatrick 1984), helper mortality may be inflated by emigration, but annual surveys of surrounding habitat patches suggest that emigration rates from our study population are low (Supplementary Methods). We used a gaussian distribution to model log-transformed fecundity and a Poisson distribution to model immigration. We modeled immigration separately for males and females to capture the impact of female-biased dispersal (Woolfenden & Fitzpatrick 1984). Please see Supplementary Methods for full details.

### 4. Contribution of Inbreeding, Density, and Environmental Variation to Population Growth

To quantify the contributions of our covariates to variation in the population growth rate, we performed a path analysis (Van Tienderen 2000) as an extension of our tLTRE (Knape *et al*. 2023). We decomposed variation in growth caused by each vital rate (quantified in our tLTRE) by the covariate associated with these rates. We analytically solved for the sensitivities of the vital rates to each environmental factor using the link functions and estimated *β* values from our GLMs. We then used these sensitivities along with the chain rule to convert the sensitivities of the population growth rate to the observed vital rates into the sensitivities of the population growth rate to each environmental factor and performed another tLTRE using the covariates in place of the vital rates. We summed the contributions to variation in the population growth rate based on each covariate. Please see Supplementary Methods for details.

### 5. Simulations

To confirm our hypothesis that changes in other vital rates compensated for changes in immigration and inbreeding, we ran post-hoc simulations to project population sizes under different scenarios. We projected our initial population vectors from 1988 using our projection matrices, assigning values for vital rates using predictions from our GLMs. This approach allows us to simulate vital rates under different conditions. We detrended vital rates by using the same value for the time fixed effect in all years. To incorporate density dependence, we recalculated population density each year based on the projected population size and used that value to predict vital rates for the next year. We simulated three scenarios: (1) decreasing immigration and increasing inbreeding with increasing breeder survival, (2) no temporal trends in breeder survival, and (3) no temporal trends in breeder survival, immigration, or inbreeding. Please see Supplemental Methods for details.

## Results

### 1. Demographic and Environmental Trends

Despite previously-documented decreases in immigration (Mann Kendall tau = -0.337, *p* = 0.007) and increases in population-wide inbreeding (Mann Kendall tau = 0.405, *p* = 0.001), the census population size remained relatively stable (coefficient of variance = 0.167; Figure 1; Reed & Hobbs 2004). In fact, we found an increase in population size over time (Mann Kendall tau = 0.273, *p* = 0.024), largely driven by an increase in the number of established pairs (Mann Kendall tau = 0.414, *p* = 0.007; Figure 1A; Table S2). Annual growth rates fluctuated rapidly between growth (*λ* > 1) and decline (*λ* < 1; Figure S1). We found no significant temporal trend in the annual growth rate (Mann Kendall tau = -0.004, *p* = 0.987), and the geometric mean of *λ* is 0.998 (Figure S1, Table S1). When we considered sex-specific vital rates, we only found a significant decline in male immigration after correcting for multiple tests (Mann Kendall tau = - 0.405, *q* = 0.010, Figure S2, Table S2). Inbreeding significantly increased over time (Figure S5, Table S3), consistent with previous work (Chen *et al*. 2016).

### 2. Contribution of Variation in Vital Rates to Population Growth

We used tLTREs to quantify how different vital rates contribute to differences in population growth over time (Figure S7). Results for our female-only and male-only models were similar, thus we report the results for the female-only model except when noted. Temporal variation in population growth rates was primarily driven by fecundity (55.1%) and survival (32.8%, Figure 2), specifically established pair fecundity (28.9%) and breeder survival (18.9%, Figure S7). Immigration accounted for 4.8% of variation in population growth rate in the female-only model and -4.3% in the male-only model due to negative covariation between male immigration and both survival and fecundity (Figure 2). Probability to pair accounted for 4.5% of the variation in the population growth rate (Figure 2).

**Figure 2.**
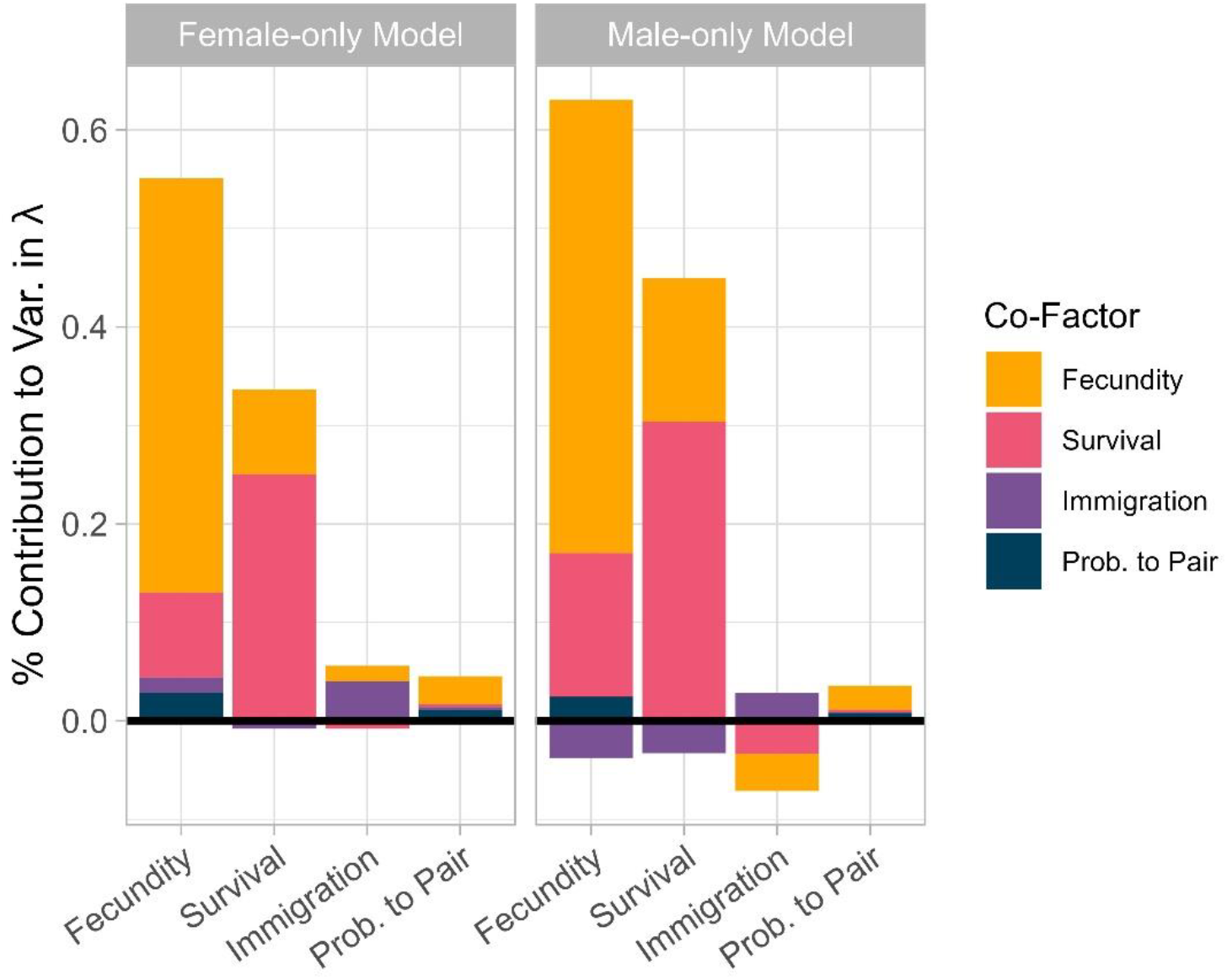
Contributions of variation in vital rates to variation in population growth rate for the female-only model (left) and male-only model (right). Bars are colored by the covariance involved (*e*.*g*., pink bars in the fecundity column are contributions made by covariance between fecundity and survival). Negative contributions indicate that the covariation between two vital rates diminished the variation in the population growth rate (*i*.*e*., one rate promoted growth while the other promoted decline).

### 3. Impact of Inbreeding, Density, and Environmental Factors on Vital Rates

To determine how inbreeding, population density, and environmental conditions influence annual variation in vital rates, we ran GLMs for each vital rate. Consistent with previous work, we found that higher inbreeding in the population was significantly associated with lower juvenile survival (*β* = -0.133, p = 0.01), male breeder survival (*β* = -0.259, p = 0.008, Table S3), and lower fecundity (*β* = -0.084, p = 0.03, Table S5). We found negative density dependence in male breeder survival (*β* = -0.170, *p* = 0.007, Table S3), female immigration (*β* = -0.166, *p* = 0.009, Table S4), both new (*β* = -0.267, *p* < 0.001) and established pair fecundity (*β* = -0.123, *p* = 0.003, Table S5), and the probability of female helpers to form new pairs (*β* = - 0.198, *p* = 0.046). Of our environmental factors, only acorn abundance had a significant association with any vital rates. Greater acorn abundance was associated with increased survival of juveniles (*β* = 0.196, *p* = 0.001), female breeders (*β* = 0.240, *p* = 0.002), and male breeders (*β* = 0.154, *p* = 0.029, Figure 3, Table S3). To account for any temporal trends that couldn’t be explained by our other fixed effects, we included a fixed effect of time. We found a significant negative effect of time on female (*β* = -0.185, *p* = 0.007) and male immigration (*β* = - 0.378, *p* < 0.001, Table S4). Though we did not find any temporal trends in breeder survival (Table S2), we found significant positive impacts of time on female (*β* = 0.338, *p* = 0.001) and male breeder (*β* = 0.417, *p* < 0.001) survival (Table S3).

**Figure 3.**
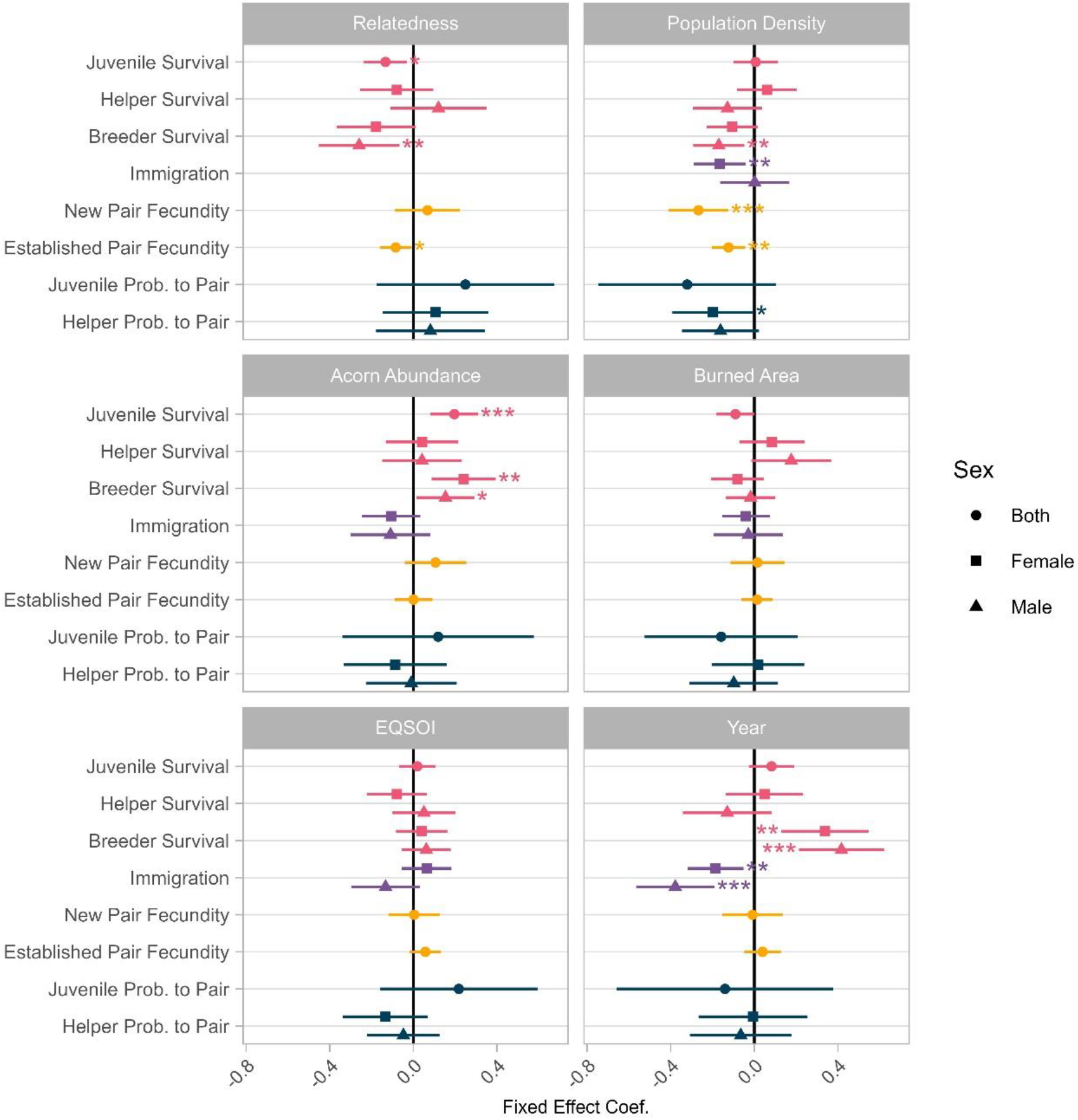
Impact of inbreeding, density, environmental variables, and year on vital rates. Error bars represent the 95% confidence interval. Colors indicate the type of vital rate modeled (pink: survival, purple: immigration, yellow: fecundity, dark blue: probability to pair). Juvenile vital rates are not divided between males and females due to the relatively large number of unsexed juveniles. Asterisks indicate significance (* *p* < 0.05, ** *p* < 0.01, *** *p* < 0.001).

### 4. Contribution of Inbreeding, Density, and Environmental Variation to Population Growth

To quantify the effect of inbreeding, density, and environmental factors on population growth, we calculated the sensitivities of population growth to these factors and summed the contribution of each factor through their impact on vital rates. Population growth rate had similar sensitivities to inbreeding, population density, acorn abundance, and temporal trends, with negative effects of inbreeding and density and positive effects of acorn abundance and time (Figure 4A). Sensitivity analysis indicates that vital rates with positive temporal trends (breeder survival) more than compensated for those with negative temporal trends (immigration) (Figure 3). When we weighted sensitivities by the observed variation in each factor, we found that variation in population density contributed the most to variation in population growth rate (12.1%), followed by variation in acorn abundance (7.1%) and EQSOI (0.6%, Figure 4B). Variation in inbreeding and temporal trends (variation in time) both contributed very little (>0.1% and 0.5%, respectively; Figure 4), largely because negative covariation between them canceled out any potential contributions to population growth (Figure 4B). Negative covariation between inbreeding and density as well as a slight decline in acorn abundance over time (Figure S5, Table S2) also reduced fluctuations in population size (Figure 4).

**Figure 4.**
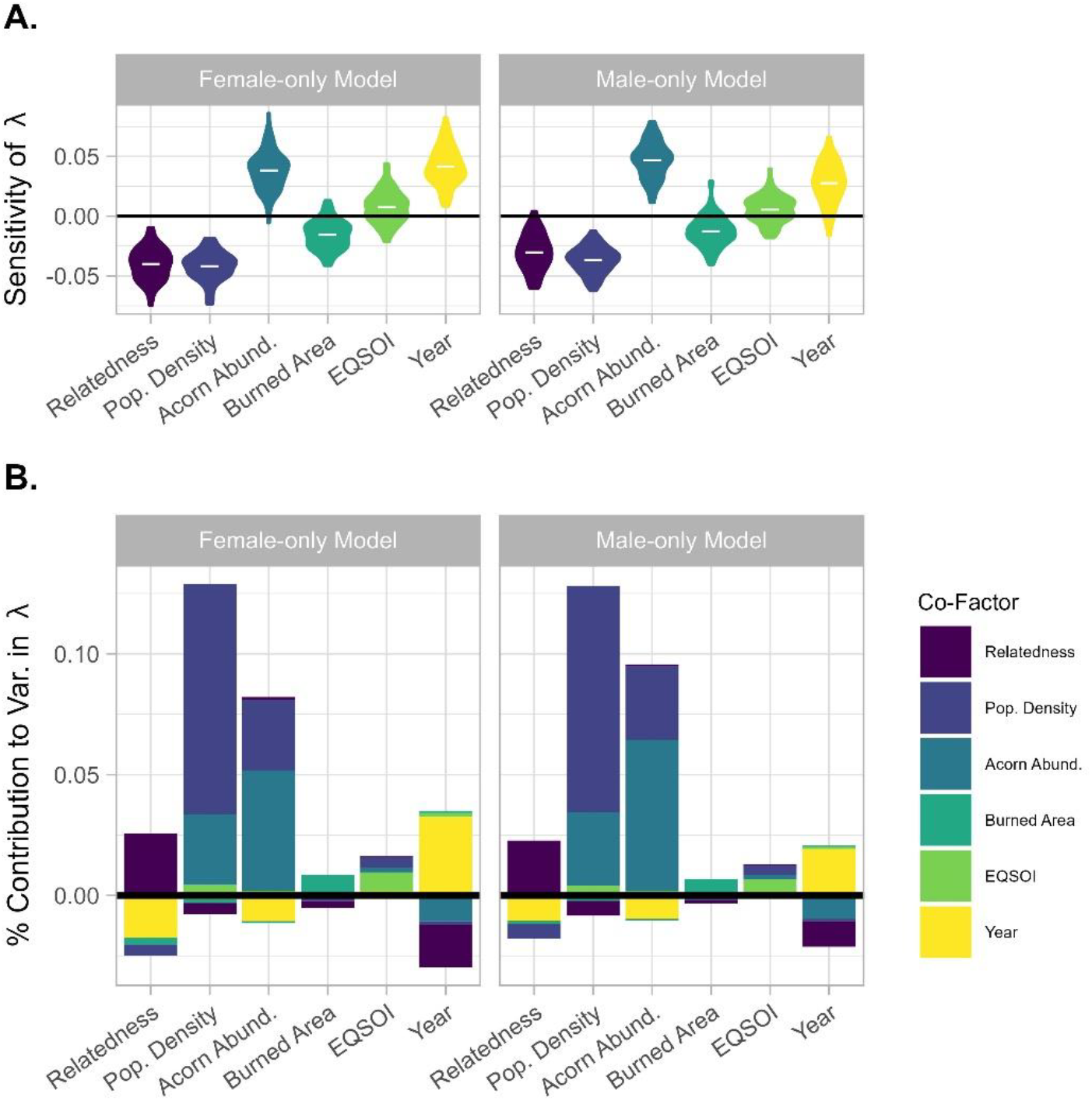
Contributions of variation in inbreeding, density, and environmental factors to variation in population growth rate. (A) Sensitivity of the population growth rate to inbreeding, density, acorn abundance, burned area, EQSOI, and year. The distributions are the result of resampling the effect sizes from the GLMs. The mean sensitivity is indicated with a white bar. (B) Inbreeding, density, and environmental factors contributed to the population growth rate through their association with vital rates. Bars are colored by covariances (*e*.*g*., the yellow bar in the acorn abundance column shows the contribution made by covariance between acorn abundance and year). Negative contributions indicate that the covariation between two factors diminished the variation in the population growth rate (*i*.*e*., one factor promoted growth while the other promoted decline).

### 5. Simulations

We simulated density-dependent population growth to test whether the observed increase in breeder survival was sufficient to compensate for the observed decline in immigration and increase in inbreeding. We first performed simulations that included all observed temporal trends as a control. While these simulations consistently underestimated population size in later years, they accurately predicted the observed population stability overall (Figure 5). Divergence between observed and simulated population dynamics is driven by a few unusually high fecundity years. When we simulated dynamics without a temporal increase in breeder survival, the population declined in both sex-specific models, with a more dramatic decrease in the male-only model (Figure 5). This decline in population size is rescued in both models when we also removed temporal trends in immigration and inbreeding (Figure 5).

**Figure 5.**
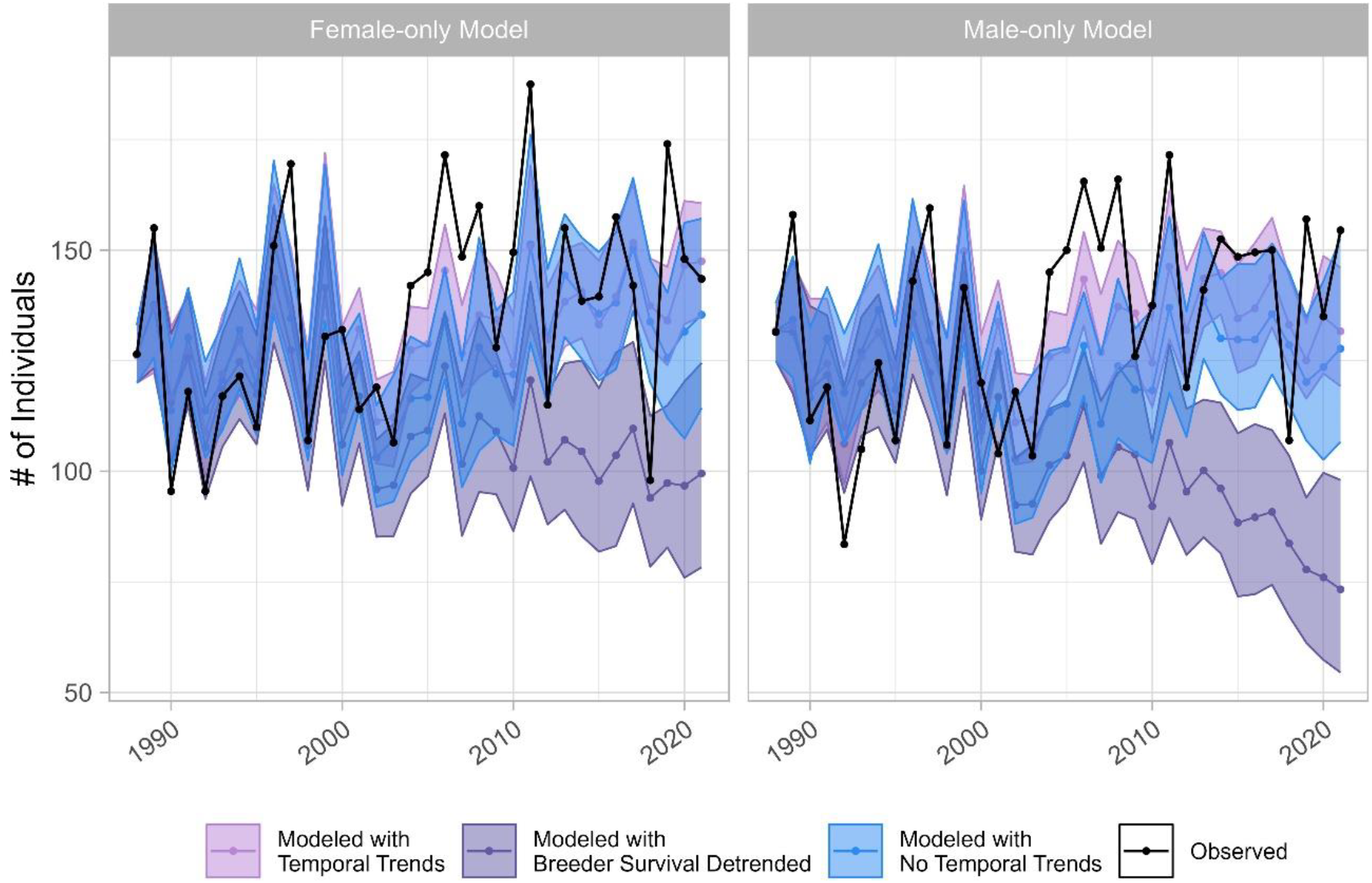
Observed and simulated population dynamics for females (left) and males (right). Density-dependent simulations that used vital rates predicted with observed temporal trends in immigration, inbreeding, and breeder survival (pink; Modeled with Temporal Trends) matched observed dynamics (black). Simulations that removed the temporal trend in breeder survival ( purple; Modeled with Breeder Survival Detrended) resulted in population decline, and we recovered observed population dynamics when we removed temporal trends in immigration, inbreeding, and breeder survival (blue; Modeled with No Temporal Trends). Ribbons indicate the 95% confidence interval for simulated population size.

## Discussion

We investigated how a wild population has maintained population size despite increasing isolation by quantifying the effect of variation in local demographic rates, immigration, inbreeding, population density, and environmental factors on variation in population growth. The stability of our Florida Scrub-Jay population results from density dependence and compensation by breeder survival. Though immigration does contribute to variability in population growth rates, we found a positive trend in breeder survival over time (after accounting for annual fluctuations in inbreeding, population density, and acorn abundance) that outweighs the demographic impact of declines in immigration and increasing inbreeding. This result contradicts our initial prediction that flexibility in the demography of non-breeders would buffer the population at saturation during changing conditions (Porter & Coulson 1987). We found direct evidence of density dependence in female immigration and covariation between male immigration and both survival and fecundity, suggesting that density dependent immigration also contributed to stability. Finally, despite evidence of inbreeding depression in vital rates with high contributions to population growth (survival and fecundity), inbreeding had a negligible effect on variation in population growth rate because of compensation by the temporal trend in breeder survival and negative covariation between inbreeding and density.

Our results demonstrate the power of long-term ecological studies to provide highly detailed insight into population regulation during changing conditions. The compensation of both declining immigration and increasing inbreeding by breeder survival, resulting in a significantly greater number of established breeders over time (Table S2), demonstrates that significant shifts in local demography were required to maintain population stability through increasing isolation. Multiple studies have documented how increasing immigration drove increases in other vital rates that allowed newly-established populations to become self-sufficient source populations (Herman & Lynch 2022; Santoro *et al*. 2016; Tenan *et al*. 2017). Our study presents a rare example of the opposite occurrence: an existing stable population adjusting to decreased immigration and becoming more self-sufficient via compensatory increases in other vital rates. Characterizing how populations can remain robust to increasing isolation is important for identifying the appropriate spatial scale for effective management of populations threatened by habitat fragmentation (Paquet *et al*. 2021)..

The observed increase in breeder survival could be a result of decreased immigration or of other trends. Because Florida Scrub-Jays engage in potentially risky territorial defense behaviors (Woolfenden & Fitzpatrick 1984), decreased immigration could have increased breeder survival through relaxed competition for breeding territories. Alternatively, immigrants frequently have reduced fitness compared to residents in birds (Barbraud & Delord 2021; Dickel *et al*. 2023), thus declining immigration could lead to increased average breeder survival. However, residents and immigrants do not have significantly different breeder lifespans in our population (Wilcoxon signed-rank test, W = 62241, *p* = 0.22). It is also possible that increased breeder survival has contributed to decreased immigration instead, via more limited breeding vacancies. The observed temporal change in breeder survival could also be caused by a reduction in negative density dependence, decreased reliance on acorns, or decreased effects of inbreeding, or as a result of successional change in the habitat due to the prescribed fire regime (Pesendorfer *et al*. 2021). While we detected no impact of burned area on breeder survival, prescribed fires can increase the availability of arthropod prey (Goud 2017) and create more optimal habitat structure that can increase breeder survival (Breininger *et al*. 2009; Fitzpatrick & Bowman 2016).

We found that inbreeding contributed relatively little to variation in the population growth rate, consistent with previous findings that inbreeding depression may not directly translate into population decline (Johnson *et al*. 2011). The overall impact of inbreeding was reduced by covariation between inbreeding and density (Figure 4B), which supports the soft selection hypothesis (Bozzuto *et al*. 2019; Keller *et al*. 2007; Wallace 1975). Populations experiencing soft selection primarily include those where intraspecific competition is important (Bell *et al*. 2019), including highly territorial species such as the Florida Scrub-Jay. Our work provides an empirical example of density dependent regulation maintaining population growth despite observable inbreeding depression. However, we note that the level of inbreeding observed in our populations remains relatively low when compared to populations experiencing extinction vortices, a negative feedback loop of population decline and inbreeding depression (Fagan & Holmes 2006; Frankham 2015; Kardos *et al*. 2023; Pimm *et al*. 2006). It remains to be seen whether demographic effects of inbreeding will change as population-wide inbreeding coefficients continue to increase. Future work should test the level of inbreeding required for a significant demographic response, with particular attention paid to effects on population density and competition for limited resources, such as breeding opportunities.

Our study emphasizes the importance of considering both males and females when modeling population dynamics when – as is often the case – sex differences exist among demographic processes. For example, we found evidence of density dependence in immigration of females but not males, which could be a result of female-biased dispersal. Similar sex-specific population dynamics have been found in sexually monomorphic species that exhibit some sex differences in vital rates (Kraus *et al*. 2008), although density dependence has been found in immigration of both sexes for other bird species that also exhibit female-biased dispersal (Wilson *et al*. 2017). Our simulations predicted greater decline without compensation by breeder survival in males compared to females, likely because of the greater decline in male immigration over time and lack of density dependent buffering in male immigration. Had we used the standard approach of only fitting female-only models, we would have underestimated the role of demographic compensation in maintaining stability and overestimated the importance of density dependent immigration.

Combining tLTREs and a path analysis is a powerful approach for evaluating the effects of inbreeding, population density, and environmental factors on population growth; however, successful application of this approach requires careful evaluation of model assumptions. Our GLMs cannot specify causation, but the tLTRE provides direct information on how observed variation in vital rates results in observed variation in population growth rate. Our retrospective analysis incorporates the contributions of covariances between vital rates or covariates to observed population dynamics, making it difficult to quantify the effects of single variables in isolation and make predictions about dynamics in unobserved conditions (an advantage of prospective analyses). We added simulations to better elucidate how changes in individual factors lead to different potential outcomes, though our simulations assume that modeled relationships between vital rates and cofactors remain consistent. Our GLMs and thus the environmental tLTRE assume a linear relationship between individual covariates and vital rates. Non-linear relationships could alter the contributions of the covariates as their values change over time, leading to over- or underestimates of the effect of that covariate on population growth depending on the functional form of the relationship. For example, Allee effects could result in a reversal of density dependence at low densities, causing the sensitivity of the population to change sign (Kramer *et al*. 2009). None of our models violated the linearity assumption (Supplementary Information). We suggest that future work consider the functional relationships between covariates and vital rates. Our matrix population models assume uniformity in vital rates between resident and immigrant individuals. This assumption does not impact predictions of population growth, but can affect estimates of the impact of immigration. We found no significant differences in vital rates between residents and immigrants in our data (Supplementary Information). If that were not the case, immigration rates could impact future vital rates and thus population growth over time. Finally, we note that uncertainty in our analyses is not randomly distributed over our study period. The proportion of unsexed individuals and pedigree incompleteness is greater during the first 10 years of our dataset (Figure 1). We do not include sex-specific vital rate estimates for juveniles (the majority of our unsexed individuals), and we found no significant difference between our pedigree-derived inbreeding coefficients for all individuals vs. a subset of individuals with more complete pedigree information (Figure S8). Future work would benefit from more precise estimates of emigration from our focal population, as density dependent emigration is another important aspect of dispersal-mediated population regulation (Doncaster *et al*. 1997; Harman *et al*. 2020). Additionally, identifying the source populations of immigrants would allow us to perform metapopulation analyses and study the impact of dispersal at different geographic scales.

We provide much needed empirical evidence on the demographic consequences of temporal variation in immigration and inbreeding. Our results show that compensation by other vital rates can maintain population growth rates as immigration declines or inbreeding increases. Our focal population thus represents a case where local density dependence is sufficient to regulate the population as it becomes more independent of the surrounding metapopulation. This result emphasizes the importance of maintaining high quality habitat that is capable of supporting such robust populations in the face of habitat fragmentation.Our study demonstrates the power of tLTREs and path analyses for characterizing potential demographic and environmental drivers of population dynamics in changing environments.

## Data Accessibility

All data and scripts used in this study are publicly available at github.com/jtsummers53/LTRE_project.

## Acknowledgments

We thank the staff and interns at Archbold Biological Station who have contributed greatly over many years to collecting the ecological and demographic data that makes this research possible. We also thank the fellow members of the Chen Lab (Kristin Hardy, Bailey Jones, Faye Romero, Shailee Shah, and Daniel Seidman) for guidance that shaped the structure of this manuscript. J.S. was supported by NIH grant 1R35GM133412 to N.C.

