## Supplementary Information for "Impacts of increasing isolation and environmental variation on Florida Scrub-Jay demography"

### SUPPLEMENTARY METHODS

#### Study Population

Researchers at Archbold Biological Station have intensively studied a population of individually-marked Florida Scrub-Jays since 1969 (Woolfenden & Fitzpatrick 1984). Nestlings are uniquely banded at 11 days of age and immigrants are banded as soon as possible upon discovery. The study site is actively managed with regular prescribed burns (Main & Menges 1997). We defined our study area using a buffer of one median territory diameter (386.5 m) around the territories monitored in 1988-1990. Individuals that do not appear in any future annual censuses are presumed dead, which includes individuals who dispersed beyond the study area. Dispersal is exceedingly rare for juveniles and breeders (Woolfenden & Fitzpatrick 1984) Annual peripheral surveys of all suitable scrub habitat tracts near Archbold Biological Station (encompassing a total area of ~700 sq. km; (Boughton & Bowman 2011; Woolfenden & Fitzpatrick 1984) indicate low emigration out of our focal population: from 29 years of surveys, we have only encountered 39 individuals that were banded as nestlings at Archbold, which comprises only ~1% of the juveniles born in our study site during that time. Immigrants are defined as individuals not present within the study area during the first year (1989) and who were not born within our study area. Prior to 1999, individuals were sexed based on behavior or breeding records, with molecular sexing for a subset of the population. After 1999, nearly all individuals were sexed molecularly (Woolfenden & Fitzpatrick 2020). Thus, nearly all (95.8%) our unsexed individuals are juveniles or helpers present in our study population before 1999. This long-term study has resulted in a multigenerational pedigree of 14,227 individuals. Our final dataset includes 4,267 individuals (1,948 female, 1,820 male, and 499 unsexed) from 1,894 territories. All work was approved by the Cornell University Institutional Animal Care and Use Committee (IACUC 2010-0015) and authorized by permits from the US Fish and Wildlife Service (TE824723-8), the US Geological Survey (banding permit 07732), and the Florida Fish and Wildlife Conservation Commission (LSSC-10-00205).

#### Estimating Vital Rates

We created a post-breeding census for our models by combining the monthly censuses between March 1st and May 30th. We assigned individuals observed on multiple territories to the territory they were observed on for the majority of the census period. For territories where a breeder disappeared during the breeding season, or there were multiple pairs (a rare occurrence), we assigned a single breeding pair based on the first pair to successfully fledge young on the territory that season. If no pairs successfully fledged young, we assigned the first pair to produce eggs, and in cases where no eggs were produced, we used the earlier recorded breeding pair. We assigned young to the socially dominant breeding pair, removing rare cases of extra pair paternity (Townsend *et al.* 2011). In Florida Scrub-Jays, fecundity depends on both individual breeding experience and pair experience (Woolfenden & Fitzpatrick 1984). The difference in fecundity between new and experienced breeding pairs (mean fecundity of new pairs = 1.41, mean fecundity of established pairs = 2.34, Wilcoxon signed-rank test V = 1220467, $p < 0.001$) is similar to the difference between new and experienced individual breeders (mean fecundity of new breeders = 1.18, mean fecundity of experienced breeders = 2.18, Wilcoxon signed-rank test V = 1646729, $p < 0.001$). We chose to separate fecundity based on pair experience rather than individual experience to avoid making assumptions about the unknown breeding history of immigrants arriving into the population. We checked for cases where individuals dispersed outside of the study area and returned at a later year (67 individuals). In each case, we added a dummy record identifying the individual as a helper to avoid doubling counting these individuals as immigrants. We used Mann Kendall tests to test for temporal trends in vital rates and Spearman's ρ to test for correlations between vital rates. We corrected for multiple tests using the Benjamini–Hochberg procedure with false discovery rate thresholds of $q$ = 0.001, 0.01, and 0.05 (Benjamini & Hochberg 1995).

**Covariates**

We calculated individual inbreeding coefficients using the "calcInbreeding" function in the *pedigree* package in R (Coster 2022) and relatedness coefficients between breeders from our multigenerational pedigree using the "kinship" function from the *kinship2* package in R (Sinnwell *et al.* 2022), setting the inbreeding coefficient of individuals with unknown parentage to 0. We used mean individual inbreeding coefficients for our survival and transition rate models and the mean relatedness coefficient of paired breeders for our fecundity models. We used GIS data of territory maps from Archbold to determine the annual occupied area for our density calculations. Because we calculated density as the number of individuals per occupied area, our density metric better captures the impact of crowding on vital rates (Rodenhouse *et al.* 1997). We acquired acorn abundance data from annual measurements of acorn production across Archbold Biological Station completed every autumn (Abrahamson & Layne 2003). We used GIS data from Archbold that maps all fires within the tract from 1967-2021 to determine the time since an area had burned. Using the *sf* (Pebesma *et al.* 2023) and *raster* (Hijmans *et al.* 2023) packages in R, we converted these maps to rasters with a resolution of 2.5x2.5m and calculated the area burned between 2-9 years ago. We downloaded monthly EQSOI measurements from the NOAA National Weather Service Center for Climate Prediction (https://www.cpc.ncep.noaa.gov/data/indices/) and averaged values across each census year.

**Stage-Structured Matrix Population Model**

As Florida Scrub-Jays are cooperative breeders with demonstrated differences in fecundity between experienced and newly-formed breeding pairs, we constructed matrix population models with four stages: juveniles, helpers, new pairs, and established pairs (Figure 1). We used the following transition matrix *A* in all analyses:

$$A = \begin{bmatrix} P_j B_j \frac{F_n}{2} & P_h B_h \frac{F_n}{2} & (P_b P_p D_n B_b + P_b(1-P_p)B_b)\frac{F_n}{2} + P_b P_p (1-D_n)\frac{F_e}{2} & (P_b P_p D_e B_b + P_b(1-P_p)B_b)\frac{F_n}{2} + P_b P_p (1-D_e)\frac{F_e}{2} \\ P_j(1-B_j) & P_h(1-B_h) & P_b P_p D_n (1-B_b) + P_b(1-P_b)(1-B_b) & P_b P_p D_e (1-B_b) + P_b(1-P_p)(1-B_b) \\ P_j B_j & P_h B_h & P_b P_p D_n B_b + P_b\left(1-P_p\right)B_b & P_b P_p D_e B_b + P_b\left(1-P_p\right)B_b \\ 0 & 0 & P_b P_p (1-D_n) & P_b P_p (1-D_e) \end{bmatrix} \quad (S1)$$

Where $P_j$, $P_h$, $P_b$, and $P_p$ are the survival rates for juveniles, helpers, breeders, and breeders' mates (note that $P_p$ is equivalent to male breeder survival in the female-only model, and female breeder survival in the male-only model);

$B_j$, $B_h$, *and* $B_b$ are the probabilities of forming new breeding pairs for juveniles, helpers, and breeders whose mates died or underwent divorce;

$F_n$ and $F_e$ are the mean number of offspring produced by new and established pairs;

*and* $D_n$ and $D_e$ are the divorce rates for new and established pairs.

Rows and columns are ordered by life stages (juveniles, helpers, new pairs, and established pairs). To include immigration in our model, we created a vector $I_m$ denoting the contribution of immigration to each life stage:

$$I_m = \left[\frac{F_n}{2}, (1 - B_I)I, B_I I, 0\right] \quad (S2)$$

Where $F_n$ is the mean number of offspring produced by new pairs;
$B_I$ is the probability of forming a new breeding pair for immigrants;
and $I$ is the number of new arriving immigrants.

We added $I_m$ to the projected population vector in order to estimate the population size in the next year:

$$N_{t+1} = A_t \times N_t + I_{m_t} \quad (S3)$$

Where $N_{t+1}$ is the population vector at time $t$ + 1;
$A_t$ is the transition matrix at time $t$;
$N_t$ is the population vector at time $t$;
and $I_{m_t}$ is the immigration vector at time $t$

Because we used a post-breeding census, individuals who were juveniles or helpers at the start of a census year have the possibility of establishing as a breeder and reproducing by the end of the census year. Thus, the fecundity of these classes is not zero, and is weighted by the probability of each becoming a breeder. The transition from a juvenile to a helper is calculated by multiplying juvenile survival ($P_j$) with the probability of *not* forming a new pair as a juvenile (1 - Bj), whereas the transition from a juvenile to a new pair is calculated by multiplying juvenile survival (Pj) with the probability of forming a new pair as a juvenile (Bj). Helper transition rates are calculated similarly. The transition from a new pair to an established pair and the persistence of an established pair requires that both members of a pair survive ($PbP_p$) and the pair does not divorce (1 - Dn or 1 - De). The transition from a new or established pair to a new pair consists of the sum of two terms: breeders who survived but lost their mate and breeders who divorced. Both terms are multiplied by the probability that the focal breeder successfully re-pairs. If the breeder does not re-pair, they would transition into a helper. We tested for independence between vital rates of different stages using a Wilcoxon signed-rank test and separated vital rates accordingly. We fit female-specific and male-specific models using sex-specific values for helper survival, breeder survival, probability of pairing for helpers, probability of re-pairing for breeders, and immigration. We decided to separate models for the two sexes for ease of interpretation, as the sensitivities of vital rates estimated from two-sex models fluctuate dramatically with small deviations from a 1:1 sex ratio (Jenouvrier *et al.* 2010). We generated population vectors for each model by summing the number of individuals assigned to each sex within each of our four stages. To account for uncertainty due to missing sex assignments, we randomly assigned sexes to unsexed individuals and performed 100 iterations. We validated our assumption of a 1:1 sex ratio for each social class (juveniles, helpers, and breeders) using $\chi^2$ tests. We found sex ratios of 0.50 for the total population ($\chi$ = 0.29, $p$ = 0.59), 0.50 for breeders ($\chi$ = 0.29, $p$ = 0.59), 0.51 for helpers ($\chi$ = 0.94, $p$ = 0.33), and 0.50 for juveniles ($\chi$ = 0.18, $p$ = 0.67).

**Transient Life Table Response Experiment**

We performed a tLTRE to quantify the contributions of different vital rates to variation in population growth rate over time while incorporating time-lagged effects. The vital rates of year $t$ - 1 can indirectly affect population growth rate through covariation with vital rates in year $t$ and directly affect population growth rate by influencing the stage structure in year $t$. The sensitivity of the growth rate to the stage structure in year *t* - 1 was near zero, which indicates that all direct effects of past vital rates are accounted for with a one year time lag. For indirect effects, we assumed that past vital rates are independent of future environment conditions and set the sensitivity of the past population growth rate on future vital rates to zero. Thus, we decomposed variation in population growth rate as:

$$Var(\lambda) = \sum_{k=0}^{1}\sum_{l=0}^{1}\sum_{i=1}^{n}\sum_{j=1}^{n} Cov\left(\theta_{i_{t-k}}, \theta_{j_{t-l}}\right)\frac{\partial\lambda}{\partial\theta_{i_{t-k}}}\frac{\partial\lambda}{\partial\theta_{j_{t-l}}} \qquad (S4)$$

Where $\theta_{i_{t-k}}$ is vital rate *i* of *n* in time period *t* - *k*,
$\theta_{j_{t-l}}$ is vital rate *j* of *n* in time period *t* - *l*,
Cov($\theta_{i_{t-k}}$, $\theta_{j_{t-l}}$) is the covariance between vital rates *i* and *j* in time periods *t* - *k* and *t* - *l*,
$\frac{\partial\lambda}{\partial\theta_{i_{t-k}}}$ is the sensitivity of $\lambda$ to $\theta_{i_{t-k}}$,
and $\frac{\partial\lambda}{\partial\theta_{j_{t-k}}}$ is the sensitivity of $\lambda$ to $\theta_{j_{t-l}}$.

The covariance between vital rates captures the variance of a single vital rate when *k* = *l* and covariation between a vital rate and its value in the following year when *k* ≠ *l*. To calculate the sensitivities used in a tLTRE, we used the mean projection matrix across all years. Thus, we averaged annual vital rates from 1988-2019 for the value of vital rates when *k* or *l* = 1, and averaged annual vital rates from 1989-2020 when *k* or *l* = 0 to calculate the sensitivity of population growth to our vital rates.

**Impact of Inbreeding, Density, and Environmental Factors on Vital Rates**

We fitted GLMs using the *glmmTMB* package (Brooks *et al.* 2023). For simplicity, we will refer to all these covariates as "environmental factors" even though we acknowledge that inbreeding and density are not pure environmental factors. Both inbreeding and population density are affected by demography, but here our focus is on the effects of inbreeding and density on demography and population dynamics. We weighted the data by the number of individuals used to calculate the annual vital rate. We used the "testResiduals" function from the package *DHARMa* (Hartig & Lohse 2022) to evaluate model residuals. We tested the uniformity of our model residuals using Kolmogorov-Smirnov tests (Berger & Zhou 2014), which compares the goodness-of-fit of fitted data to observed values. We also compared the dispersion of fitted and observed values to ensure that the assumed variances of our model match the observed data. The juvenile survival model contained 6 outlier data points that caused the residuals of the model to violate our assumptions. We excluded these years (1989, 1993, 1996, 1997, 2006, and 2019) from the final model. We used the package *performance* (Lüdecke *et al.* 2023) to check for multicollinearity among covariates using variable inflation factors (VIFs). None of our fixed effects presented a concern (VIF < 5 for all, Table S7). We conducted all analyses in R version 4.3.1 (R Core Team 2021).

**Sensitivity of Vital Rates to Environmental Factors**

We derived the sensitivities of the vital rates to our environmental factors by taking the partial derivatives of the link functions for each GLM with respect to the environmental factors. For our logistic models (survival and probability to pair) we used the logit link function. The partial derivative with respect to environmental factor $x_i$ is therefore:

$$\frac{\partial P}{\partial x_i} = \frac{\beta_i e^{\beta_0+\beta_1 x_1+\cdots\beta_n x_n}}{(e^{\beta_0+\beta_1 x_1+\cdots\beta_n x_n}+1)^2} \qquad (S5)$$

Where *P* is the probability of an event occurring,

$\beta_n$ is the effect estimate for environmental factor *n*,
and $x_n$ is the mean value of environmental factor *n.*

For our Poisson model (immigration) and log-transformed Gaussian model (fecundity), we used the log link function. The partial derivative with respect to environmental factor $x_i$ is therefore:

$$\frac{\partial \mu}{\partial x_i} = \beta_i e^{\beta_0 + \beta_1 x_1 + \cdots \beta_n x_n} \qquad (S6)$$

Where $\mu$ is the mean number of events that occur,
$\beta_n$ is the effect estimate for environmental factor *n*,
and $x_n$ is the mean of the annual values of the environmental factor *n*.

**Environmental Transient Life Table Response Experiment**

We further partitioned the contributions of the vital rates by environmental factors using the chain rule to determine the sensitivity of the growth rate to each environmental factor ($e_h$, for all environmental factors *h*):

$$\frac{\partial \lambda}{\partial e_{h_{t-k}}} = \sum_{i=1}^{n} \frac{\partial \lambda}{\partial \theta_{i_{t-k}}} \frac{\partial \theta_{i_{t-k}}}{\partial e_{h_{t-k}}} \qquad (S7)$$

where $\frac{\partial \lambda}{\partial e_{h_{t-k}}}$ is the sensitivity of $\lambda$ to $e_h$ in time period *t - k*,
$\frac{\partial \lambda}{\partial \theta_{i_{t-k}}}$ is the sensitivity of $\lambda$ to $\theta_i$ in time period *t - k,*
and $\frac{\partial \theta_{i_{t-k}}}{\partial e_{h_{t-k}}}$ is the sensitivity of vital rate $\theta_i$ to $e_h$ in time period *t - k.*

We used the partial derivatives of the link functions from the GLMs to calculate the sensitivity of each vital rate to each environmental factor ($\frac{\partial \theta_{i_{t-k}}}{\partial e_{h_{t-k}}}$, Equation S5 and S6) using effect sizes estimated from the GLMs. To account for uncertainty in our models, we randomly selected 100 effect size values from a normal distribution with mean and variance obtained from our GLMs. We calculated the sensitivity of the vital rates to the environmental factors for two time periods (1990-2020 and 1989-2019) to create sensitivities for both the current and time-lagged vital rates in the tLTRE. We adapted Equation S4 to quantify how variation in the environmental variables contributed to variation in population growth rate through their impacts on the vital rates:

$$Var(\lambda) = \sum_{k=0}^{1} \sum_{h=1}^{n} \sum_{m=1}^{n} Cov\left(e_{h_{t-k}}, e_{m_{t-k}}\right) \frac{\partial \lambda}{\partial e_{h_{t-k}}} \frac{\partial \lambda}{\partial e_{m_{t-k}}} \qquad (S8)$$

where $e_{h_{t-k}}$ is environmental factor *h* of *n* in time period *t - k*,
$e_{m_{t-k}}$ is environmental factor *m* of *n* in time period *t - k*,
$Cov(e_{h_{t-k}}, e_{m_{t-k}})$ is the covariance between environmental factors *h* and *m* within time periods *k* and *l*,
$\frac{\partial \lambda}{\partial e_{h_{t-k}}}$ is the sensitivity of $\lambda$ to $e_{h_{t-k}}$,
and $\frac{\partial \lambda}{\partial e_{m_{t-k}}}$ is the sensitivity of $\lambda$ to $e_{m_{t-k}}$.
We report the mean contribution of each environmental factor in the main text.

**Simulations**

To confirm our hypothesis that changes in other vital rates compensated for changes in immigration and inbreeding, we ran post-hoc simulations projecting population size changes over our study period under different scenarios. We started the simulations with the observed sex-specific population vectors from 1988 (randomly assigning sexes for unsexed individuals for each iteration, as before and predicted the population vector of the next year by multiplying these vectors by our transition matrix $A(t)$ and adding the immigration vector $I_m(t)$. We used our GLMs to predict new values for the vital rates used in $A(t)$ and $I_m(t)$. We randomly selected each value from a normal distribution of predicted values with mean and variance obtained from our GLMs to account for uncertainty in our models. To simulate different conditions, we adjusted the fixed effect values of the input data when predicting vital rates with our GLMs. To remove temporal trends from immigration and breeder survival, we set the time fixed effect to the earliest value. For inbreeding, we set each inbreeding metric to its minimum observed value. To incorporate density dependence, we calculated the population density resulting from each annual projection of the population vector and used that value in our vital rate predictions. All density calculations assumed that the population occupied the same area as the observed data for each year. For each sex-specific model, we simulated three scenarios: (1) a control scenario with observed temporal trends of decreasing immigration, increasing inbreeding, and increasing breeder survival; (2) a scenario that eliminated temporal trends in breeder survival; and (3) a scenario that eliminated temporal trends in breeder survival, immigration, and inbreeding. The covariates within the control simulations do not differ from the observed data, except that density is recalculated after projecting each year. Each sex-specific simulation was repeated 500 times. We compared projected population dynamics to the observed sex-specific population sizes over time. We split the number of unsexed individuals evenly between the sexes, which is a reasonable assumption given the 1:1 sex ratio in our study population.

### SUPPLEMENTARY RESULTS AND DISCUSSION

#### Correlations among vital rates and GLM covariates

We found relatively high correlations between survival rates of different stages, with significant correlations between juvenile survival and both breeder (Spearman's $\rho = 0.58$) and helper survival (Spearman's $\rho = 0.5$; Figure S3) after correcting for multiple tests. Fecundity of new and established pairs was significantly correlated (Spearman's $\rho = 0.6$), as was the probability of helpers forming pairs and breeders re-forming pairs (Spearman's $\rho = 0.5$; Figure S3). Between different vital rates, the only significant correlations were between the probability of pairing in juveniles and immigration (Spearman's $\rho = 0.57$) as well as the probability of pairing in helpers and established pair fecundity (Spearman's $\rho = 0.5$; Figure S3). The correlations within survival and fecundity likely indicate shared beneficial conditions across life stages, whereas the correlations between probabilities to pair and both immigration and fecundity indicate shared density dependence. Across our female-only and male-only models, nearly all significant correlations were between the sex-specific versions of the same vital rates (Figure S4). The only exceptions were significant cross-sex correlations between new and established pair fecundity, which are identical between the sex-specific models, and a correlation between established pair fecundity and the probability of female helpers to form pairs (Spearman's $\rho = 0.53$; Figure S4). Beyond a significant negative correlation between acorn abundance and time (Spearman's $\rho = -0.42$), the only other significant correlations between GLM covariates were across inbreeding metrics (Figure S6).

We also compared vital rates between immigrants and residents to determine whether changes in immigration may influence vital rates. We compared helper and breeder survival, helper probability to pair, breeder probability to pair, and divorce and fecundity rates for both new and established pairs using Wilcoxon rank sum tests. After adjusting for multiple tests, we found no significant differences between vital rates calculated from individuals who immigrated to the study tract and from those who were born on the study tract (all $q > 0.2$).

#### Verifying inbreeding coefficients

To ensure that differences in pedigree completeness do not bias our inbreeding estimates, we calculated the annual mean inbreeding coefficients for individuals with known family histories going back two generations (*i.e.*, individuals for whom we know all four grandparents). We found no significant difference between inbreeding coefficients calculated from the full dataset and those calculated from the subset of individuals with known ancestry (Wilcoxon signed-rank test V = 530, $p = 0.563$; Figure S8).

#### Simulations

We simulated density-dependent growth under three scenarios: (1) observed temporal trends in breeder survival, immigration, and inbreeding; (2) no temporal trends in breeder survival; and (3) no temporal trends in breeder survival, immigration, or inbreeding. The geometric mean of the growth rate for both the female and male-specific simulations with no changes in temporal trends was 1, with no variation across replicates. Detrending breeder survival caused the geometric mean of the growth rate to decrease to 0.99 (95% CI = 0.994-0.986) in the female-only simulation and to 0.982 (95% CI = 0.987-0.976) in the male-only simulation. Further removing temporal trends in immigration and inbreeding rebounded the geometric mean of the growth rate to 1 (95% CI = 1-0.997) in the female-only simulation and 0.998 (95% CI = 1-0.996) in the male only simulation. These results support our conclusion that population stability is maintained through temporal trends in breeder survival that counterbalance temporal trends in immigration and inbreeding.

#### Effects of fecundity and fire on Florida Scrub-Jay demography

Our results showing the importance of breeder survival and environmental fluctuations on population dynamics are largely consistent with previous work on the Florida Scrub-Jay (Breininger *et al.* 2022, 2023; Breininger & Oddy 2004; Woolfenden & Fitzpatrick 1984). The large contributions of variation in fecundity and population density both result from high temporal autocorrelation in new pair fecundity (Spearman's $\rho = -0.36$, $p = 0.04$), with high fecundity years frequently followed by low fecundity years (Woolfenden & Fitzpatrick 1984). Unlike other studies (Breininger 2009; Breininger *et al.* 2022, 2023; Woolfenden & Fitzpatrick 1984), we did not find an association between fire history and variation in vital rates at the population-wide scale. This discrepancy is likely due to the low variation in the proportion of burned area observed across our study area during the study period. The current burn regime was started in 1981 (Main & Menges 1997), and the amount of burned scrub across the study area as a whole has been more or less constant. However, post-fire succession and changing climate have caused declines in acorn abundance at Archbold (Pesendorfer *et al.* 2021). Thus, a population-wide interaction between fire and climate has occurred, but is acting through an average decline in acorn abundance.

**SUPPLEMENTARY FIGURES**

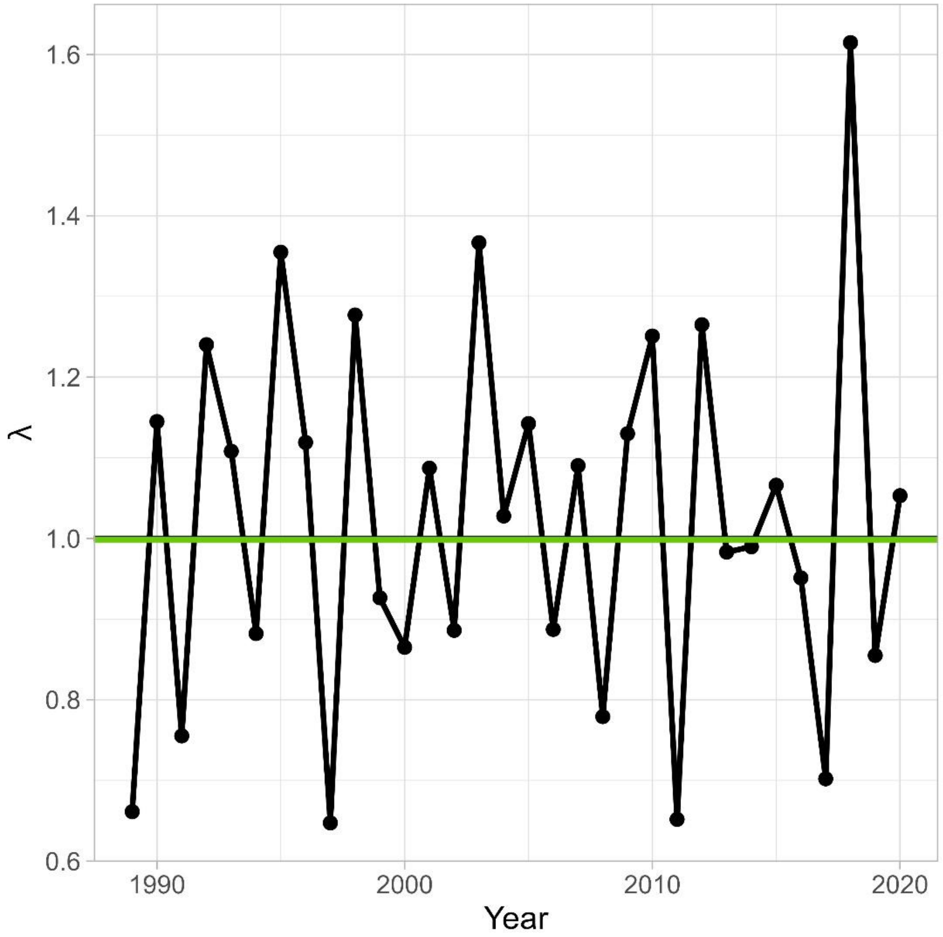


**Figure S1.** Observed annual population growth rate (λ). The geometric mean of the population growth rate (0.998) is indicated by the green line.

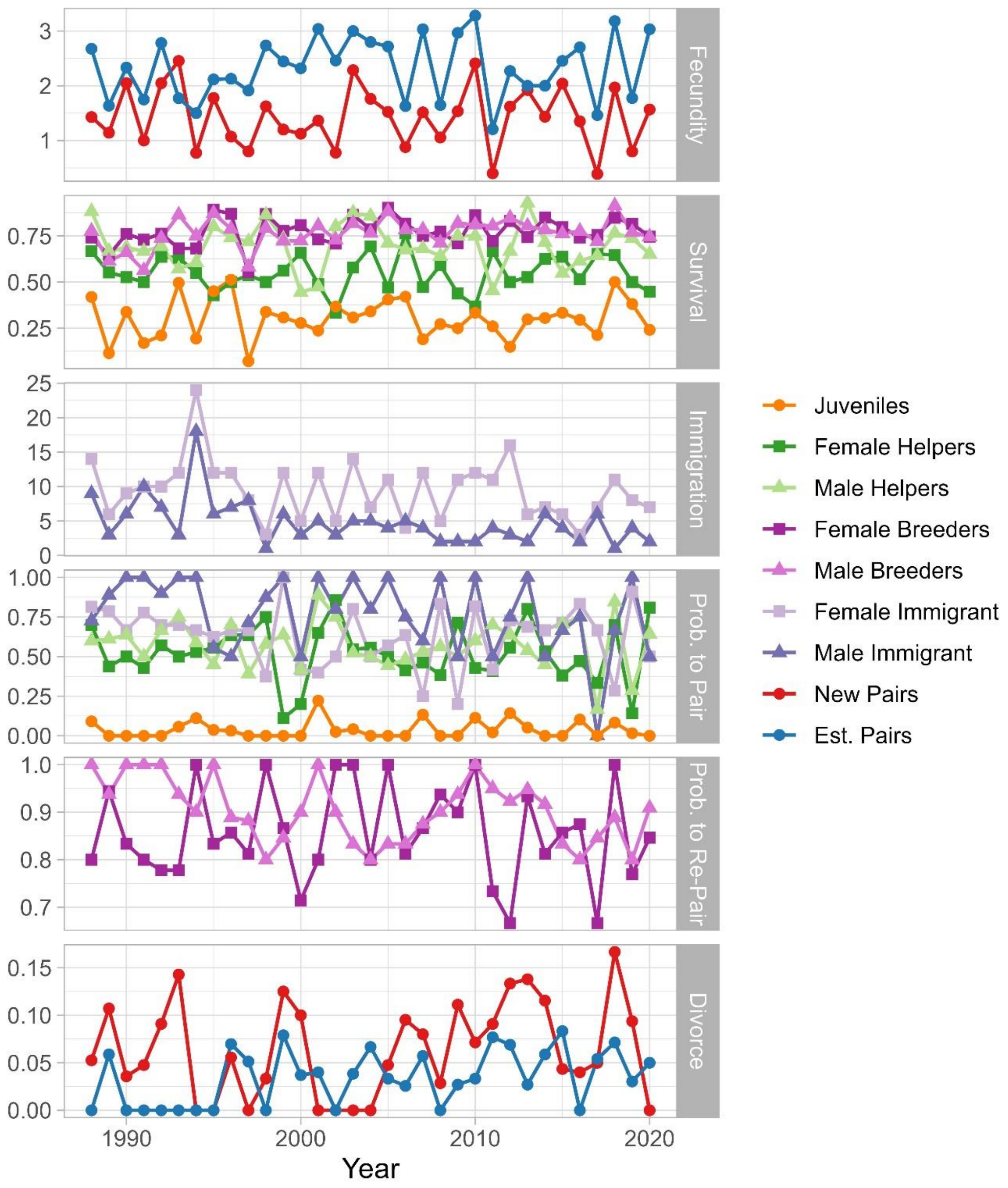


**Figure S2.** Observed vital rates (fecundity, survival, immigration, probability of pairing or re-pairing, and divorce) based on data for our study population between 1988-2021. We present sex-specific values for adult survival, immigration, and probability of pairing. Fecundity and divorce rates differ between new and established breeding pairs. We used data from 1988 to calculate the time-lagged effect of previous vital rates for the first year in the transient life table response experiments.

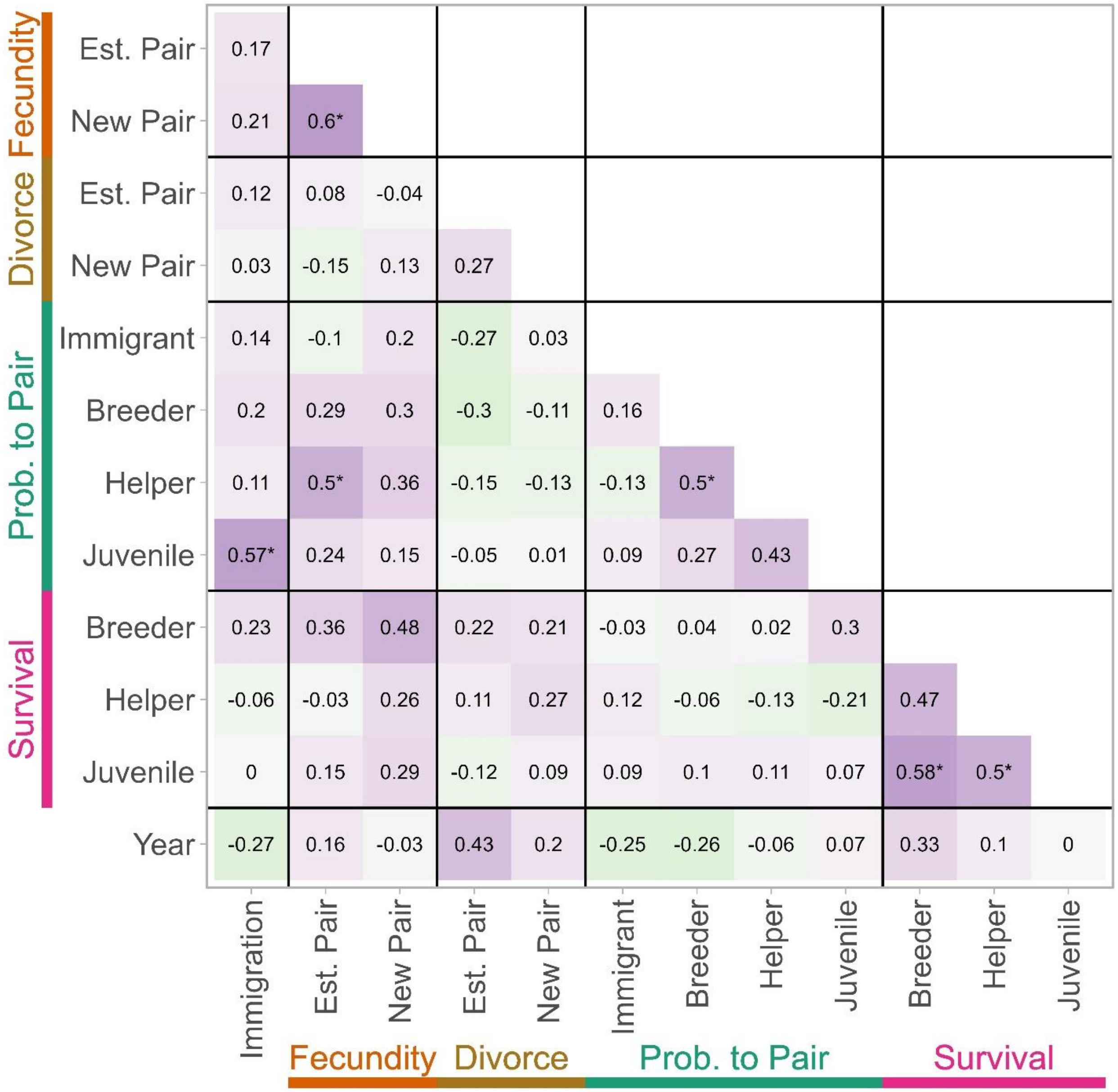


**Figure S3.** Correlations between all observed vital rates and time. Purple indicates positive correlations and green negative correlations. Values are Spearman's rho, and asterisks indicate significance after adjusting for multiple tests (* $q < 0.05$, ** $q < 0.01$, *** $q < 0.001$).

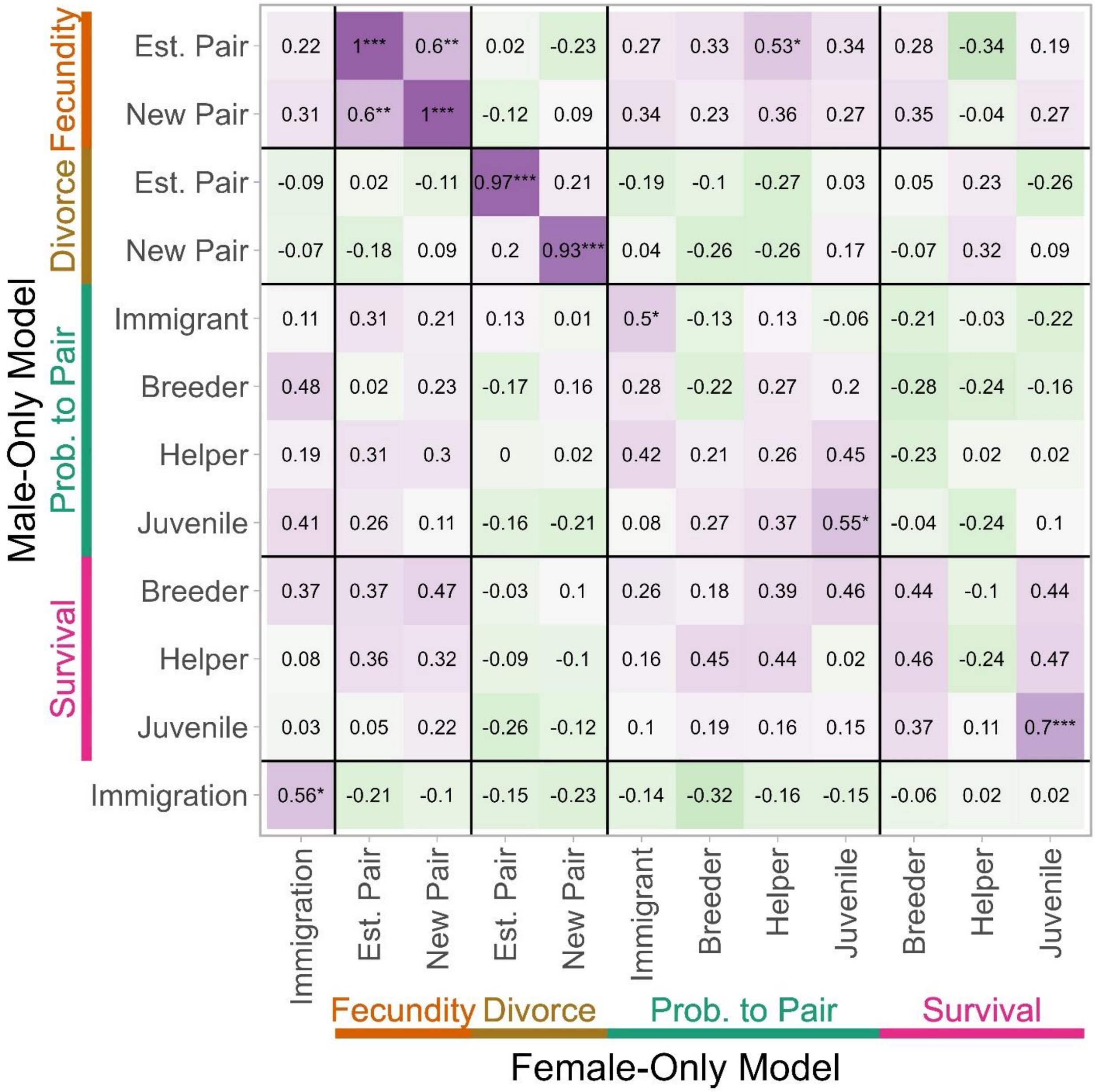


**Figure S4.** Correlations between female-only and male-only vital rate estimates. Values are Spearman's rho, and asterisks indicate significance after adjusting for multiple tests (* $q < 0.05$, ** $q < 0.01$, *** $q < 0.001$).

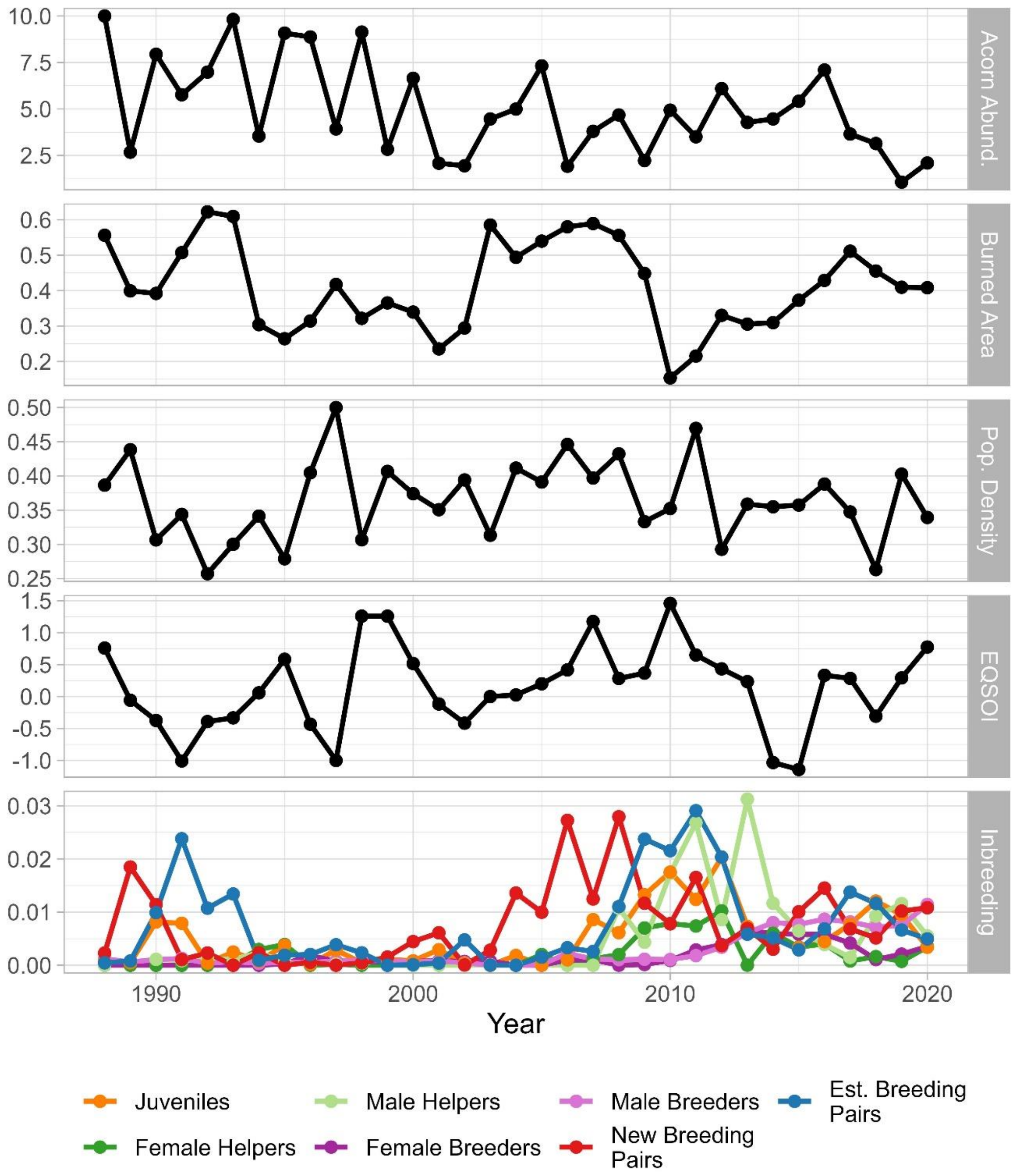


**Figure S5.** Observed variation in environmental factors between 1988-2020. We measured the mean number of acorns per tree stand (acorn abundance), the proportion of total area burned (burned area), the number of individuals per hectare (population density), EQSOI, and the mean individual inbreeding coefficient for juveniles, helpers, and breeders or the relatedness coefficient of breeders in a pair.

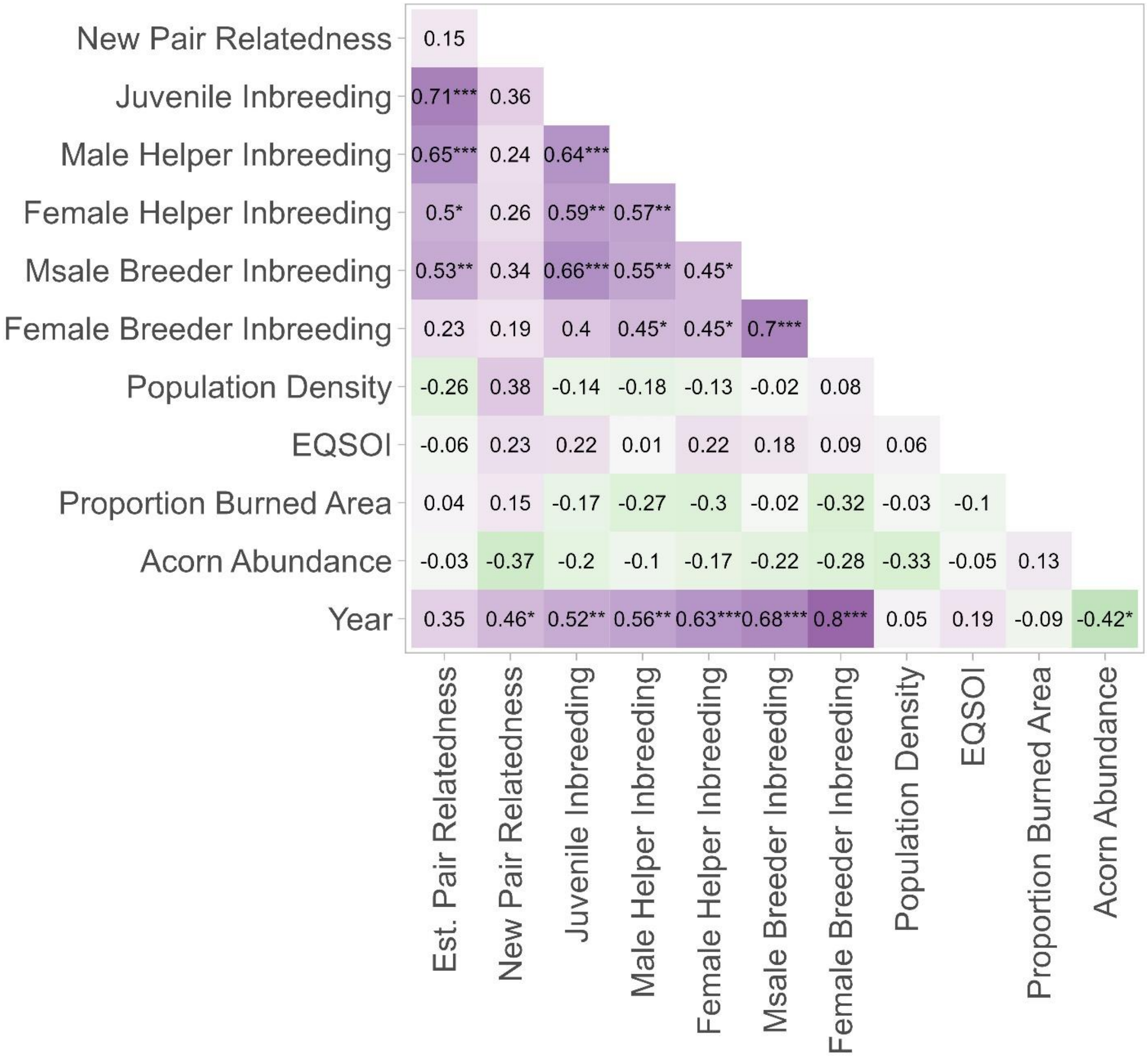


**Figure S6.** Correlations between environmental factors. Values are Spearman's $\rho$, and asterisks indicate significance after adjusting for multiple tests (* $q < 0.05$, ** $q < 0.01$, *** $q < 0.001$).

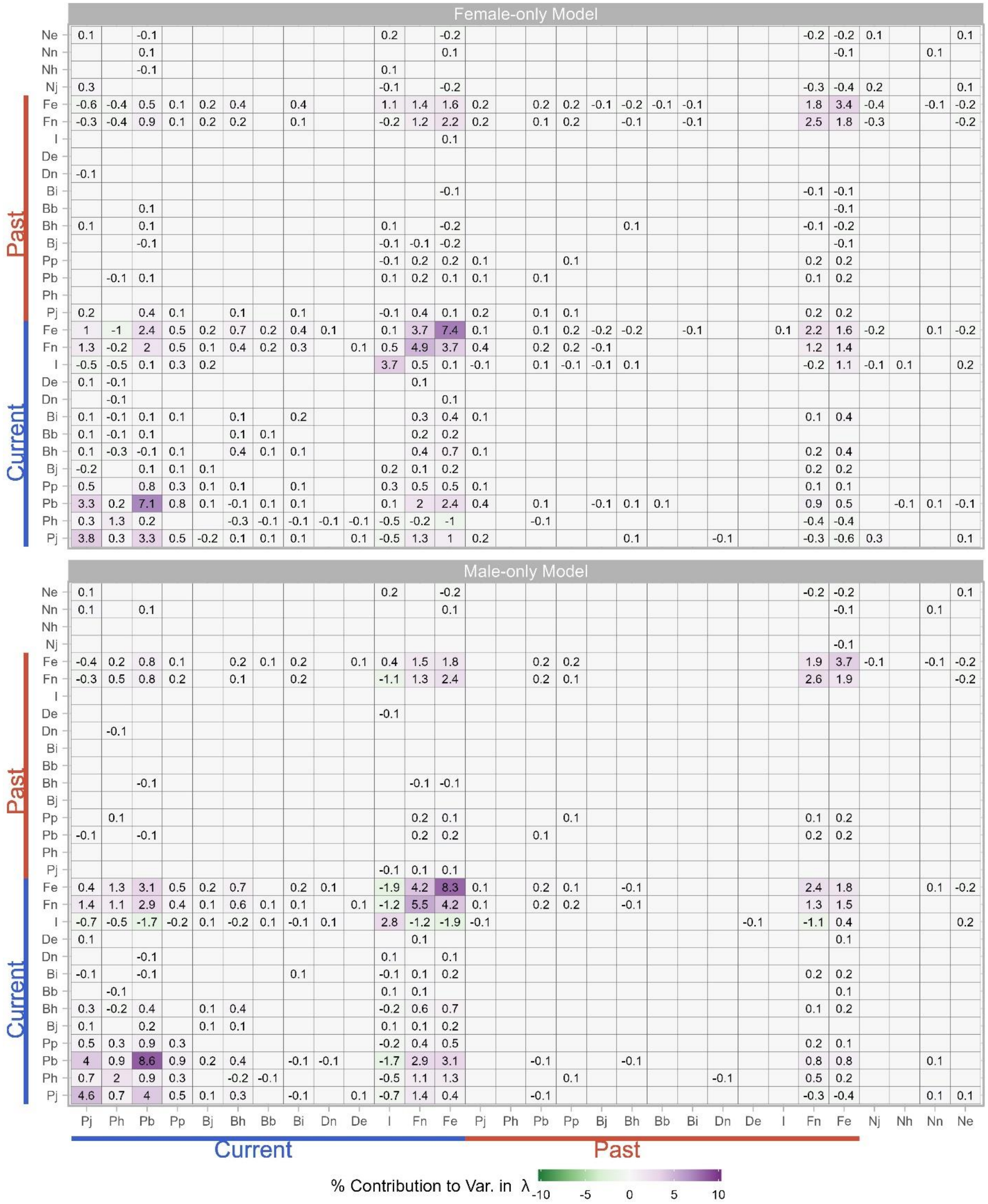


**Figure S7.** Contributions of variation and covariation in vital rates as well as the population stage structure to observed variation in population growth rate. We considered vital rates from both the current year (blue) and the previous year (red), with the previous year vital rates contributing to variation in the population growth rate through their impact on demographic structure. The vital rates are: the survival of juveniles, helpers, breeders, and breeders' mates ($P_j$, $P_h$, $P_b$, $P_p$); the probability to form a new pair for juveniles, helpers, breeders, and immigrants ($B_j$, $B_h$, $B_b$, $B_i$); the divorce rates of new and established pairs ($D_n$, $D_e$); the immigration rate ($I$); and the fecundity of new and established pairs ($F_n$, $F_e$). The population stage structure of the previous year is the proportion of the population that were juveniles, helpers, new pairs, and established pairs ($N_j$, $N_h$, $N_n$, $N_e$). We show nonzero mean contributions across 100 replicates with randomly assigned sex for unsexed individuals.

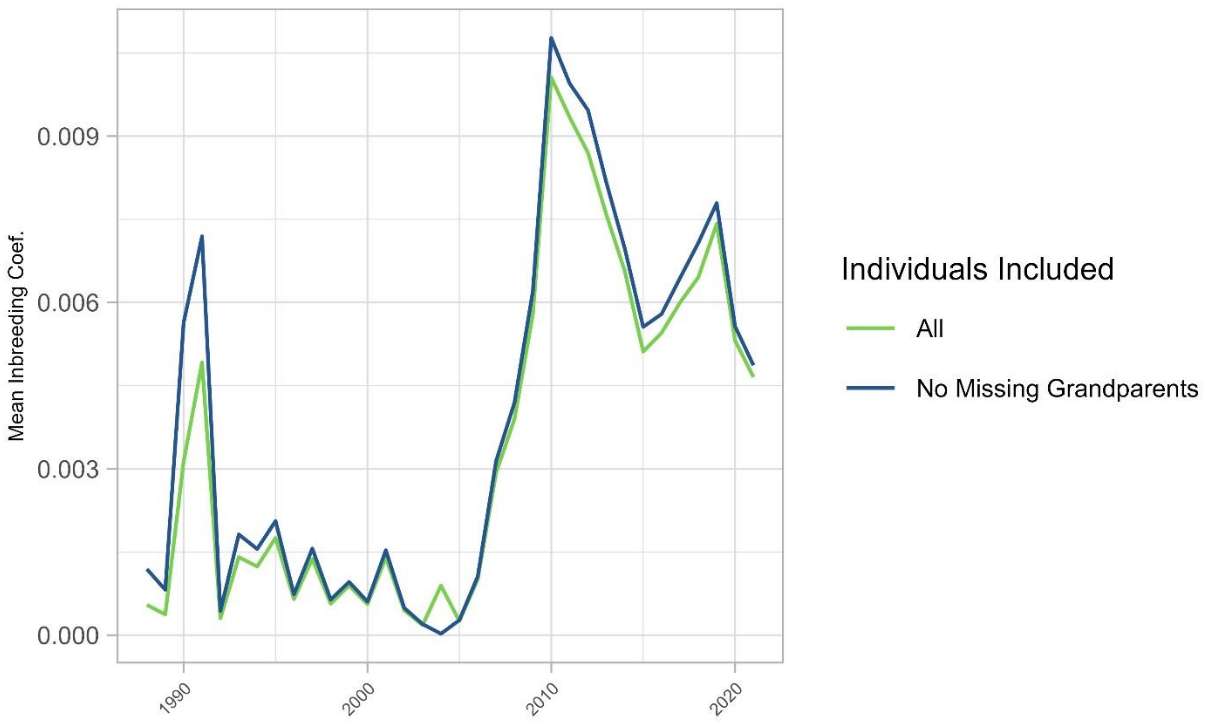


**Figure S8.** Comparison of population-wide average inbreeding coefficients over time calculated with all individuals (light green) or with only individuals with no missing grandparents (dark green). Missing grandparents are the result of incomplete pedigree records, we may underestimate the inbreeding coefficient of individuals with missing grandparents.

| Model | Geometric Mean Growth Rate | Mean Population Size |
|---|---|---|
| Both-sex Observations | 0.998 (0.23) | 270.2 (45.8) |
| Female-only Model | 1.002 (0.239) | 135.6 (23.8) |
| Male-only Model | 0.997 (0.227) | 134.2 (22.3) |

**Table S1.** Mean population growth rate and annual size. We present average values with standard deviation in parentheses calculated from direct observation of all individuals (both-sex observations) as well as from female-only and male-only models with randomly-assigned sexes for unsexed individuals. Note that the population size estimates for the female-only and male-oly models are only for one sex, which is approximately half the total population size (the sex ratio of our study population is close to 1:1).

| Category | Factor | Tau | p-value | q-value |
|---|---|---:|---:|---:|
| Environmental Factors | Acorn Abundance | -0.295 | 0.016 | 0.060 |
| | Burned Area | -0.061 | 0.631 | 0.833 |
| | Population Density | 0.038 | 0.768 | 0.915 |
| | EQSOI | 0.140 | 0.258 | 0.448 |
| Inbreeding | Juveniles | **0.330** | **0.007** | **0.040** |
| | Female Helpers | **0.446** | **0.000** | **0.005** |
| | Male Helpers | 0.306 | 0.014 | 0.058 |
| | Female Breeders | **0.612** | **0.000** | **0.000** |
| | Male Breeders | **0.495** | **0.000** | **0.001** |
| | New Pairs | 0.313 | 0.011 | 0.052 |
| | Est. Pairs | 0.237 | 0.055 | 0.150 |
| Survival | Juvenile | 0.000 | 1.000 | 1.000 |
| | Female Helper | -0.033 | 0.804 | 0.915 |
| | Male Helper | -0.051 | 0.687 | 0.872 |
| | Female Breeder | 0.140 | 0.258 | 0.448 |
| | Male Breeder | 0.166 | 0.182 | 0.367 |
| Probability to Pair | Juvenile | 0.035 | 0.802 | 0.915 |
| | Female Helper | -0.071 | 0.577 | 0.805 |
| | Male Helper | -0.015 | 0.913 | 0.973 |
| | Female Breeder | 0.010 | 0.950 | 0.980 |
| | Male Breeder | -0.289 | 0.023 | 0.077 |
| | Female Immigrant | -0.091 | 0.474 | 0.711 |
| | Male Immigrant | -0.232 | 0.076 | 0.188 |
| Divorce | New Pair | 0.133 | 0.295 | 0.464 |
| | Established Pair | 0.285 | 0.026 | 0.077 |
| Fecundity | New Pair | -0.015 | 0.914 | 0.973 |

| Category | Factor | Tau | p-value | q-value |
|---|---|---|---|---|
| | Established Pair | 0.131 | 0.292 | 0.464 |
| Immigration | Female Immigration | -0.168 | 0.189 | 0.367 |
| | Male Immigration | **-0.405** | **0.002** | **0.010** |
| Stage Distribution | Juvenile | 0.183 | 0.141 | 0.310 |
| | Helper | 0.218 | 0.080 | 0.188 |
| | New Pair | -0.070 | 0.586 | 0.805 |
| | Established Pair | **0.396** | **0.001** | **0.010** |

**Table S2.** Results of Mann Kendall tests for temporal trends in environmental factors, mean levels of inbreeding, vital rates, and the population stage distribution (*i.e.*, the number of individuals in each life stage). Significant results using an FDR threshold of 0.05 are indicated in bold.

| | Juvenile Survival | Female Helper Survival | Male Helper Survival | Female Breeder Survival | Male Breeder Survival |
|---|---|---|---|---|---|
| Intercept | -0.885*** | 0.110 | 0.818*** | 1.282*** | 1.216*** |
| | (0.044) | (0.070) | (0.079) | (0.058) | (0.057) |
| Acorn Abundance | 0.196*** | 0.042 | 0.042 | 0.241** | 0.154* |
| | (0.058) | (0.088) | (0.097) | (0.078) | (0.070) |
| Burned Area | -0.090 | 0.084 | 0.177 | -0.081 | -0.017 |
| | (0.046) | (0.080) | (0.097) | (0.065) | (0.060) |
| Time | 0.083 | 0.049 | -0.129 | 0.338** | 0.417*** |
| | (0.055) | (0.094) | (0.108) | (0.106) | (0.104) |
| EQSOI | 0.019 | -0.079 | 0.051 | 0.040 | 0.062 |
| | (0.045) | (0.073) | (0.077) | (0.063) | (0.060) |
| Population Density | 0.007 | 0.061 | -0.128 | -0.106 | -0.170** |
| | (0.054) | (0.073) | (0.084) | (0.062) | (0.062) |
| Inbreeding Coefficient | -0.133* | -0.079 | 0.120 | -0.179 | -0.259** |
| | (0.053) | (0.089) | (0.118) | (0.097) | (0.099) |
| Num.Obs. | 27 | 33 | 33 | 33 | 33 |
| R2 Marg. | 0.013 | 0.009 | 0.018 | 0.026 | 0.031 |
| AIC | 195.3 | 164.8 | 164.8 | 179.6 | 170.9 |
| BIC | 204.4 | 175.2 | 175.3 | 190.1 | 181.4 |
| RMSE | 0.07 | 0.09 | 0.11 | 0.05 | 0.05 |

* p < 0.05, ** p < 0.01, *** p < 0.001

**Table S3.** Results of the GLMs for juvenile, helper, and breeder survival. We include the standard error in parentheses.

| | Female Immigration | Male Immigration |
|---|---|---|
| Intercept | 2.225*** | 1.495*** |
| | (0.058) | (0.085) |
| Acorn Abundance | -0.106 | -0.109 |
| | (0.071) | (0.098) |
| Burned Area | -0.040 | -0.028 |
| | (0.058) | (0.085) |
| Time | -0.185** | -0.378*** |
| | (0.068) | (0.096) |
| EQSOI | 0.064 | -0.133 |
| | (0.060) | (0.084) |
| Population Density | -0.166** | 0.002 |
| | (0.064) | (0.084) |
| Num.Obs. | 33 | 33 |
| AIC | 190.6 | 156.6 |
| BIC | 199.6 | 165.5 |
| RMSE | 3.74 | 2.66 |

* p < 0.05, ** p < 0.01, *** p < 0.001

**Table S4.** Results of the GLMs for female and male immigration. We include the standard error in parentheses. We show the standard deviation of the estimated random effect intercepts.

| | | New Pair Fecundity | Est. Pair Fecundity |
|---|---|---|---|
| | Intercept | 0.262*** | 0.810*** |
| | | (0.059) | (0.037) |
| | Acorn Abundance | 0.106 | 0.001 |
| | | (0.075) | (0.046) |
| | Burned Area | 0.015 | 0.014 |
| | | (0.066) | (0.038) |
| | Time | -0.008 | 0.040 |
| | | (0.073) | (0.045) |
| | EQSOI | 0.004 | 0.058 |
| | | (0.063) | (0.038) |
| | Population Density | -0.267*** | -0.123** |
| | | (0.073) | (0.041) |
| | Pair Relatedness | 0.068 | -0.084* |
| | | (0.079) | (0.039) |
| Dispersion | Intercept | 0.339 | 0.211 |
| | Num.Obs. | 33 | 33 |
| | R2 Marg. | 0.080 | 0.022 |
| | AIC | 55.5 | 60.5 |
| | BIC | 67.5 | 72.5 |
| | RMSE | 0.34 | 0.21 |

* p < 0.05, ** p < 0.01, *** p < 0.001

**Table S5.** Results of the GLMs for fecundity. We include the standard error in parentheses. We show the standard deviation of the estimated random effect intercepts.

| | Juvenile Prob. to Pair | Female Helper Prob. to Pair | Male Helper Prob. to Pair |
|---|---|---|---|
| Intercept | -3.430*** | 0.281** | 0.322*** |
| | (0.188) | (0.097) | (0.088) |
| Acorn Abundance | 0.119 | -0.087 | -0.010 |
| | (0.234) | (0.126) | (0.111) |
| Burned Area | -0.158 | 0.019 | -0.098 |
| | (0.187) | (0.112) | (0.108) |
| Time | -0.140 | -0.006 | -0.065 |
| | (0.264) | (0.133) | (0.124) |
| EQSOI | 0.218 | -0.133 | -0.048 |
| | (0.193) | (0.104) | (0.088) |
| Population Density | -0.321 | -0.198* | -0.162 |
| | (0.217) | (0.099) | (0.094) |
| Inbreeding Coefficient | 0.249 | 0.107 | 0.082 |
| | (0.216) | (0.129) | (0.133) |
| Num.Obs. | 33 | 33 | 33 |
| R2 Marg. | 0.090 | 0.020 | 0.013 |
| AIC | 106.8 | 162.0 | 163.1 |
| BIC | 117.3 | 172.5 | 173.6 |
| RMSE | 0.05 | 0.19 | 0.15 |

* p < 0.05, ** p < 0.01, *** p < 0.001

**Table S6.** Results for the GLMs for probability of forming a new pair. We include the standard error in parentheses.

| Model | Acorn Abundance | Proportion Burned Area | Year | EQSOI | Population Density | Inbreeding Coef./Relatedness |
|---|---|---|---|---|---|---|
| Juvenile Survival | 1.37 | 1.24 | 1.45 | 1.16 | 1.15 | 1.44 |
| Female Helper Survival | 1.41 | 1.25 | 1.71 | 1.06 | 1.17 | 1.71 |
| Male Helper Survival | 1.37 | 1.42 | 1.86 | 1.02 | 1.23 | 1.86 |
| Female Breeder Survival | 1.77 | 1.21 | 3.47 | 1.25 | 1.21 | 3.05 |
| Male Breeder Survival | 1.53 | 1.02 | 3.52 | 1.15 | 1.25 | 3.05 |
| Juvenile Prob. to Pair | 2.12 | 1.51 | 2.28 | 1.36 | 1.31 | 2.09 |
| Female Helper Prob. to Pair | 1.46 | 1.28 | 1.75 | 1.04 | 1.18 | 1.72 |
| Male Helper Prob. to Pair | 1.50 | 1.37 | 1.93 | 1.02 | 1.21 | 1.79 |
| New Pair Fecundity) | 1.57 | 1.21 | 1.50 | 1.10 | 1.47 | 1.75 |
| Est. Pair Fecundity | 1.54 | 1.05 | 1.44 | 1.04 | 1.20 | 1.09 |
| Male Immigration | 1.53 | 1.07 | 1.31 | 1.07 | 1.22 | |
| Female Immigration | 1.61 | 1.07 | 1.40 | 1.09 | 1.21 | |

**Table S7.** Variance Inflation Factors (VIFs) for all fixed effects used in our GLMs.

| | Survival | | | | | Prob. to Pair | | | | | | | Divorce | | Fecundity | | Immigration | |
|---|---|---|---|---|---|---|---|---|---|---|---|---|---|---|---|---|---|---|
| Year | Juvenile | Female Helper | Male Helper | Female Breeder | Male Breeder | Juvenile | Female Helper | Male Helper | Female Breeder | Male Breeder | Female Immigrant | Male Immigrant | New Pair | Est. Pair | New Pair | Est. Pair | Female | Male |
| 1988 | 0.42 | 0.67 | 0.88 | 0.74 | 0.78 | 0.09 | 0.70 | 0.60 | 0.80 | 1.00 | 0.81 | 0.73 | 0.05 | 0.00 | 1.43 | 2.68 | 14 | 9 |
| 1989 | 0.11 | 0.55 | 0.67 | 0.65 | 0.61 | 0.00 | 0.44 | 0.61 | 0.94 | 0.94 | 0.79 | 0.89 | 0.11 | 0.06 | 1.14 | 1.64 | 6 | 3 |
| 1990 | 0.34 | 0.53 | 0.69 | 0.76 | 0.66 | 0.00 | 0.50 | 0.64 | 0.83 | 1.00 | 0.67 | 1.00 | 0.04 | 0.00 | 2.05 | 2.33 | 9 | 6 |
| 1991 | 0.17 | 0.50 | 0.67 | 0.73 | 0.56 | 0.00 | 0.43 | 0.50 | 0.80 | 1.00 | 0.78 | 1.00 | 0.05 | 0.00 | 1.00 | 1.75 | 10 | 10 |
| 1992 | 0.21 | 0.64 | 0.69 | 0.76 | 0.74 | 0.00 | 0.57 | 0.67 | 0.78 | 1.00 | 0.70 | 0.90 | 0.09 | 0.00 | 2.05 | 2.78 | 10 | 7 |
| 1993 | 0.50 | 0.63 | 0.57 | 0.68 | 0.86 | 0.06 | 0.50 | 0.75 | 0.78 | 0.94 | 0.70 | 1.00 | 0.14 | 0.00 | 2.45 | 1.77 | 12 | 3 |
| 1994 | 0.19 | 0.55 | 0.61 | 0.68 | 0.75 | 0.11 | 0.53 | 0.59 | 1.00 | 0.90 | 0.67 | 1.00 | 0.00 | 0.00 | 0.77 | 1.50 | 24 | 18 |
| 1995 | 0.45 | 0.43 | 0.80 | 0.89 | 0.87 | 0.04 | 0.56 | 0.45 | 0.83 | 1.00 | 0.62 | 0.56 | 0.00 | 0.00 | 1.78 | 2.12 | 12 | 6 |
| 1996 | 0.51 | 0.50 | 0.74 | 0.87 | 0.79 | 0.03 | 0.64 | 0.70 | 0.86 | 0.89 | 0.67 | 0.50 | 0.06 | 0.07 | 1.07 | 2.13 | 12 | 7 |
| 1997 | 0.07 | 0.54 | 0.72 | 0.55 | 0.58 | 0.00 | 0.64 | 0.39 | 0.81 | 0.88 | 0.67 | 0.71 | 0.00 | 0.05 | 0.80 | 1.91 | 8 | 8 |
| 1998 | 0.34 | 0.50 | 0.86 | 0.87 | 0.79 | 0.00 | 0.75 | 0.58 | 1.00 | 0.80 | 0.38 | 0.88 | 0.03 | 0.00 | 1.62 | 2.74 | 3 | 1 |
| 1999 | 0.31 | 0.56 | 0.73 | 0.78 | 0.72 | 0.00 | 0.11 | 0.64 | 0.87 | 0.85 | 1.00 | 1.00 | 0.12 | 0.08 | 1.20 | 2.44 | 12 | 6 |
| 2000 | 0.28 | 0.66 | 0.44 | 0.81 | 0.72 | 0.00 | 0.20 | 0.42 | 0.71 | 0.90 | 0.42 | 0.50 | 0.10 | 0.04 | 1.12 | 2.32 | 5 | 3 |
| 2001 | 0.24 | 0.49 | 0.47 | 0.73 | 0.80 | 0.22 | 0.65 | 0.89 | 0.80 | 1.00 | 0.40 | 1.00 | 0.00 | 0.04 | 1.36 | 3.04 | 12 | 5 |
| 2002 | 0.37 | 0.33 | 0.80 | 0.71 | 0.73 | 0.03 | 0.86 | 0.75 | 1.00 | 0.90 | 0.50 | 0.80 | 0.00 | 0.00 | 0.78 | 2.46 | 5 | 3 |
| 2003 | 0.31 | 0.58 | 0.88 | 0.86 | 0.82 | 0.04 | 0.55 | 0.52 | 1.00 | 0.83 | 0.80 | 1.00 | 0.00 | 0.04 | 2.29 | 3.00 | 14 | 5 |
| 2004 | 0.34 | 0.69 | 0.86 | 0.78 | 0.76 | 0.00 | 0.56 | 0.50 | 0.80 | 0.80 | 0.50 | 0.80 | 0.00 | 0.07 | 1.76 | 2.80 | 7 | 5 |
| 2005 | 0.40 | 0.47 | 0.71 | 0.90 | 0.88 | 0.00 | 0.50 | 0.44 | 1.00 | 0.83 | 0.57 | 1.00 | 0.05 | 0.03 | 1.52 | 2.72 | 11 | 4 |
| 2006 | 0.42 | 0.76 | 0.68 | 0.82 | 0.78 | 0.00 | 0.41 | 0.48 | 0.81 | 0.83 | 0.64 | 0.75 | 0.10 | 0.03 | 0.88 | 1.63 | 4 | 5 |
| 2007 | 0.19 | 0.47 | 0.68 | 0.75 | 0.78 | 0.13 | 0.46 | 0.53 | 0.87 | 0.88 | 0.25 | 0.60 | 0.08 | 0.06 | 1.51 | 3.03 | 12 | 4 |
| 2008 | 0.27 | 0.59 | 0.64 | 0.77 | 0.71 | 0.00 | 0.38 | 0.56 | 0.94 | 0.90 | 0.83 | 1.00 | 0.03 | 0.00 | 1.06 | 1.65 | 5 | 2 |
| 2009 | 0.25 | 0.44 | 0.75 | 0.71 | 0.82 | 0.00 | 0.71 | 0.50 | 0.90 | 0.94 | 0.20 | 0.50 | 0.11 | 0.03 | 1.54 | 2.97 | 11 | 2 |
| 2010 | 0.33 | 0.37 | 0.75 | 0.86 | 0.81 | 0.11 | 0.43 | 0.60 | 1.00 | 1.00 | 0.82 | 1.00 | 0.07 | 0.03 | 2.41 | 3.28 | 12 | 2 |
| 2011 | 0.26 | 0.67 | 0.45 | 0.72 | 0.80 | 0.02 | 0.41 | 0.70 | 0.73 | 0.95 | 0.42 | 0.50 | 0.09 | 0.08 | 0.40 | 1.21 | 11 | 4 |
| 2012 | 0.15 | 0.50 | 0.67 | 0.83 | 0.85 | 0.14 | 0.56 | 0.64 | 0.67 | 0.92 | 0.73 | 0.75 | 0.13 | 0.07 | 1.62 | 2.27 | 16 | 3 |
| 2013 | 0.30 | 0.53 | 0.93 | 0.74 | 0.80 | 0.05 | 0.80 | 0.54 | 0.93 | 0.95 | 0.69 | 1.00 | 0.14 | 0.03 | 1.92 | 2.00 | 6 | 2 |
| 2014 | 0.31 | 0.62 | 0.71 | 0.85 | 0.78 | 0.00 | 0.53 | 0.45 | 0.81 | 0.92 | 0.67 | 0.50 | 0.12 | 0.06 | 1.43 | 2.00 | 7 | 6 |
| 2015 | 0.33 | 0.64 | 0.55 | 0.80 | 0.76 | 0.00 | 0.38 | 0.71 | 0.86 | 0.83 | 0.71 | 0.67 | 0.04 | 0.08 | 2.04 | 2.45 | 6 | 4 |
| 2016 | 0.30 | 0.52 | 0.62 | 0.74 | 0.78 | 0.10 | 0.47 | 0.75 | 0.88 | 0.80 | 0.83 | 0.75 | 0.04 | 0.00 | 1.35 | 2.70 | 3 | 2 |
| 2017 | 0.21 | 0.65 | 0.64 | 0.75 | 0.72 | 0.00 | 0.33 | 0.17 | 0.67 | 0.85 | 0.67 | 0.00 | 0.05 | 0.05 | 0.39 | 1.46 | 7 | 6 |
| 2018 | 0.50 | 0.65 | 0.76 | 0.85 | 0.91 | 0.08 | 0.70 | 0.85 | 1.00 | 0.89 | 0.29 | 0.67 | 0.17 | 0.07 | 1.97 | 3.18 | 11 | 1 |
| 2019 | 0.38 | 0.50 | 0.74 | 0.82 | 0.77 | 0.02 | 0.14 | 0.29 | 0.77 | 0.80 | 0.91 | 1.00 | 0.09 | 0.03 | 0.80 | 1.77 | 8 | 4 |
| 2020 | 0.24 | 0.45 | 0.65 | 0.75 | 0.75 | 0.00 | 0.81 | 0.64 | 0.85 | 0.91 | 0.50 | 0.50 | 0.00 | 0.05 | 1.57 | 3.03 | 7 | 2 |

**Table S8.** Observed vital rates from 1988-2020 calculated using raw census data for males and females. We did not calculate sex-specific vital rates for juveniles because there were too many unsexed juveniles.
